# Astrocytes instructively regulate neuronal translation

**DOI:** 10.64898/2026.07.27.741020

**Authors:** Wendy J Liu, Carson C Schultz, Eshaan A Khan, Mauricio M Oliveira, Srinidhi V Kalavai, Charles J Sheehan, Xunzhao Zhang, Richard Sam, Eric Klann

## Abstract

Neuronal protein synthesis is essential for synaptic plasticity and long-term memory, yet whether its regulation is shaped by other cell types remains poorly understood. Here, we show that astrocyte-secreted proteins regulate global neuronal translation depending on astrocytic state. Astrocyte-conditioned medium (ACM) increased neuronal translation under basal conditions, an effect enhanced by astrocyte stimulation with the activity-dependent factor BDNF, whereas ACM from neurotoxic reactive astrocytes, a state linked to neuroinflammation and Alzheimer’s disease, suppressed neuronal translation. Across these conditions, neuronal mTORC1 activity consistently tracked with translational output, whereas the integrated stress response (ISR) acted through distinct, state-specific mechanisms that did not always track with neuronal translation. Furthermore, we identified astrocyte-secreted apolipoprotein E (APOE) and its associated cargo as a negative regulator of neuronal translation that contributed to the decreased translation induced by neurotoxic reactive astrocytes. We also found that astrocyte-secreted signals required neuronal endocytosis to influence translation and drove synaptic remodeling dependent on glutamatergic signaling and neuronal mTORC1 activity. Together, these findings identify astrocytes as active, instructive regulators of neuronal translation and synaptic structure, with implications for understanding how astrocyte dysfunction may disrupt the translational mechanisms underlying impairments in synaptic plasticity and long-term memory in neurodegenerative disease.

**GRAPHICAL ABSTRACT:** 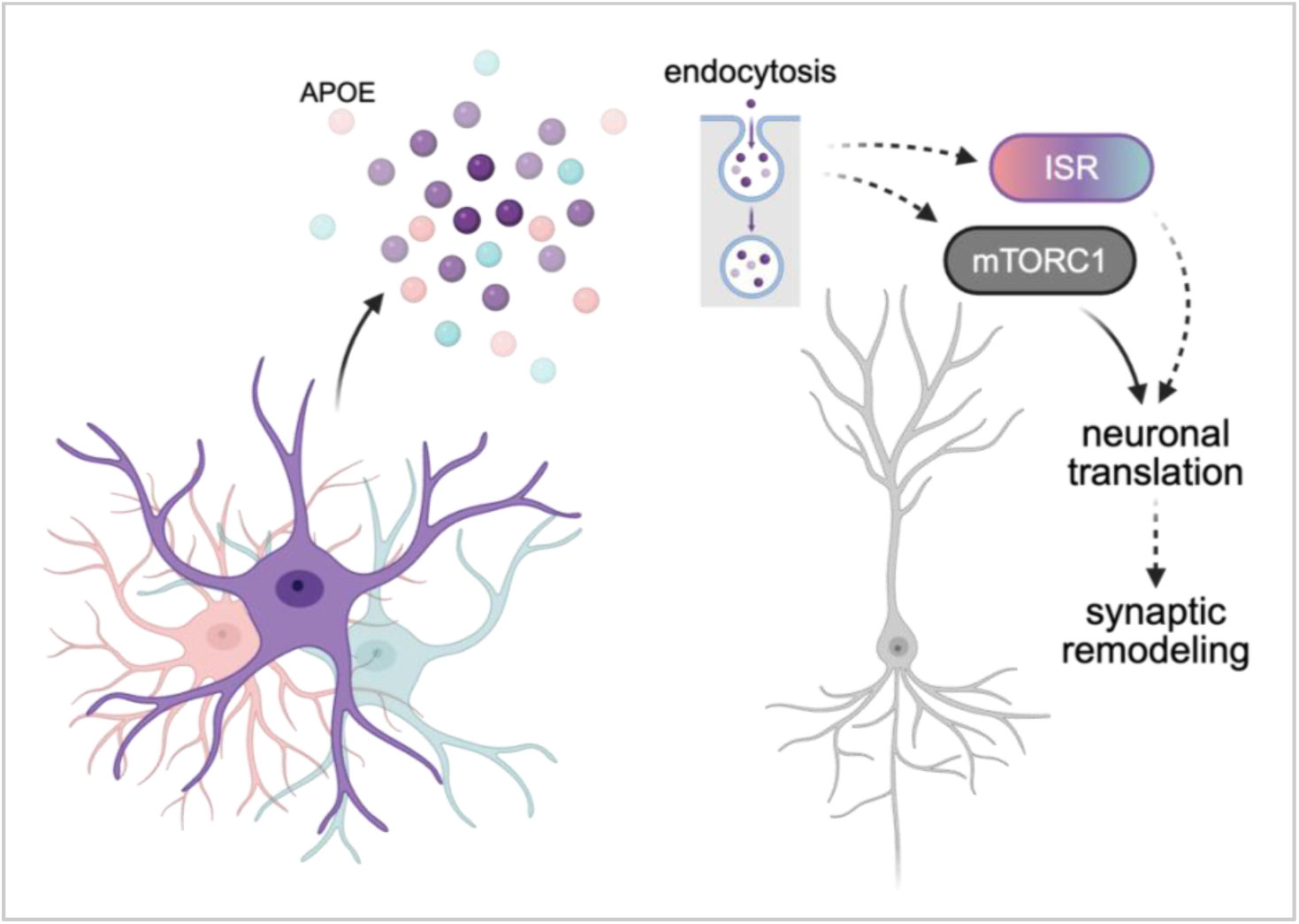

## Introduction

Protein synthesis, or translation, is a highly regulated process integral for the formation of long-lasting synaptic plasticity and long-term memory^1–4^. Over the past six decades, advances in molecular and genetic technologies have propelled extensive characterization of translation regulation, and how each component of its regulatory pathways is critical for plasticity and memory.

The most complex and highly regulated stage of translation is initiation. Over twenty-five different initiation factors in eukaryotes (eIFs) are involved in coordinating ribosome subunits and mRNA sequences^5^, and this coordination is further regulated by two major signaling pathways, 1) mammalian target of rapamycin complex 1 (mTORC1) signaling and 2) the integrated stress response (ISR)^3,4^. mTORC1 activity can release inhibitors of eIFs that recognize the 5’ mRNA cap to promote initiation, through various kinases and substrates. On the other hand, kinases and effectors of the ISR center around the phosphorylation of eIF2α (p-eIF2α), which inhibits general translation initiation. p-eIF2α can also stimulate translation of specific factors including growth arrest and DNA damage-inducible protein 34 (GADD34), a scaffolding protein that recruits protein phosphatase 1 (PP1) to dephosphorylate p-eIF2α, thereby relieving translational repression and serving as negative feedback. The intricacy of these multilayered regulatory networks highlights how maintaining translation at optimal levels is critical for physiological function.

The high degree of complexity of translational control suggests that in addition to the fine-tuning of protein synthesis that occurs in response to physiological activity, vulnerability to severe imbalances could contribute to disease. For example, in healthy contexts, activating mTORC1 signaling or disrupting ISR activity to increase global translation usually results in lower thresholds for inducing synaptic plasticity and promotes memory, whereas altering activity in the opposite direction to lower translation results in impaired plasticity and memory^6–8^. The latter phenomenon appears to be more prevalent in neurodegenerative disorders such as Alzheimer’s disease (AD): dysregulated levels of various mTORC1 and ISR components are observed in postmortem AD human brains^6,9–18^, and pharmacological or genetic manipulation of this signaling to increase translation initiation can improve AD-related plasticity and memory impairments in AD model mice^16,19–21^. Despite this progress, however, the mechanisms underlying translational dysregulation in AD remain incompletely understood, in part because the broader cellular context in which this dysregulation occurs has been largely overlooked.

Across AD and physiological contexts, studies characterizing translation and its regulation in the nervous system have focused almost exclusively on neurons: not only are they the “electrical wires” of the brain, but they also exhibit long-lasting plasticity hypothesized to be the basis of memory. Despite limited prior investigation^22,23^, the extent to which the regulation of neuronal translation is cell-autonomous, or is instead actively shaped by other cell types, remains largely unexplored. Astrocytes are among the most abundant non-neuronal cell types in the brain, and secrete a wide range of trophic and supporting factors that promote neuronal survival and modulate synaptic structure and function^24–27^. Rather than providing only passive support, research from the past two decades has demonstrated that astrocytes actively participate in synaptic activity. Structurally, fine astrocyte processes ensheathe synapses and can either increase or decrease their coverage in response to synaptic activity, leading to increased spine stabilization^28,29^. Functionally, astrocytes can uptake and respond to neurotransmitters via increasing cytosolic Ca^2+^, and release biologically active molecules such as glutamate, GABA, D-serine, ATP, and multiple peptides that can regulate synaptic transmission^30–34^. At the cognitive level, astrocyte activity is important for multiple functions, including memory^34–39^.

Beyond these physiological contributions to synaptic structure, function, and cognition, astrocytes also play a role in neurodegenerative diseases including AD^40–43^. Many physiological functions of astrocytes, including calcium signaling, glutamate uptake, and amyloid-beta clearance, are dysregulated or lost in AD^40^. In addition to the loss of homeostatic support usually provided by astrocytes during neuroinflammation, which is observed in AD, astrocytes will instead secrete pro-inflammatory and toxic factors that decrease synapse number and strength, and can eventually lead to neuronal degeneration and death^22,41^. Collectively, these cellular and molecular disruptions are likely to have profound consequences on the synaptic and circuit-level function that underlies cognition. Yet despite the clear involvement of astrocytes in both healthy and disease-state brain function, it remains unknown how homeostatic or reactive astrocyte states directly contribute to plasticity and memory.

Given this dual role of astrocytes, supporting neuronal function in physiological conditions and contributing to dysfunction in disease, as well as the established importance of translation in plasticity and memory, we hypothesized that astrocytes secrete molecules that regulate neuronal translation, and that this regulation may change depending on the astrocytic state. To examine this question, we leveraged a simple model using primary neural cell cultures, which have the advantage of isolating secreted factors in the form of conditioned cell culture medium. We demonstrate that astrocyte-secreted proteins regulate neuronal translation and the mTORC1 and ISR translational control pathways. Furthermore, this regulation changes in response to inflammation or to BDNF activity and is important for synaptic plasticity. We also reveal that astrocyte-secreted proteins regulate neuronal translation via an endocytic mechanism, and that translational impairments caused by inflammation can be rescued by targeting multiple points in this pathway, including astrocyte protein synthesis, a specific secreted factor, and neuronal endocytosis. Collectively, our findings reveal a novel astrocyte-to-neuron mode of communication that may be important for not only physiological nervous system function, but also pathology in neurodegenerative diseases.

## Results

### Astrocytes secrete proteins that regulate neuronal translation

To determine whether astrocytes influence neuronal translation, we generated primary mouse astrocyte cultures and collected astrocyte conditioned medium (ACM), which was then applied to primary mouse neurons. Global translation in these neurons was assessed one of two ways: 1) transient labeling with puromycin, an aminonucleoside that is incorporated into nascent polypeptides which can then be detected with anti-puromycin antibodies^44^, or 2) labeling with l-azidohomoalanine (AHA), a noncanonical methionine analog that contains an azide moiety which, after incorporating into nascent polypeptides, can be detected using click chemistry to fluorescent ligands^45^. Compared to non-treated controls, neurons treated with ACM displayed increased translation (Fig 1B-C). Notably, different lengths of astrocyte conditioning time resulted in similar increases in neuronal translation (Fig S1). Given that culture medium collected after shorter astrocyte conditioning intervals may contain higher concentrations of metabolites, which are also known to be important for neuronal translation and memory^46,47^, Fig 1C includes data for only the longest condition time, 72h, and this is the timepoint we used moving forward. Consistent with increased global translation, neurons treated with ACM also displayed increased phosphorylation of ribosomal protein S6 (p-S6) (Fig 1C, right), consistent with elevated mTORC1 signaling. On the other hand, while we predicted ACM-treated neurons to display decreased ISR signaling, we observed decreased GADD34-eIF2α interaction without a corresponding change in total GADD34 levels (Fig 1D-E, S2), suggesting that this change in interaction occurs independent of GADD34 abundance. These results indicate that ACM-induced translation is accompanied by increased mTORC1 signaling, whereas the effects of ACM on GADD34 activity deviate from our initial prediction, a pattern we will later examine further across different astrocyte states and stimuli.

**Figure 1.**
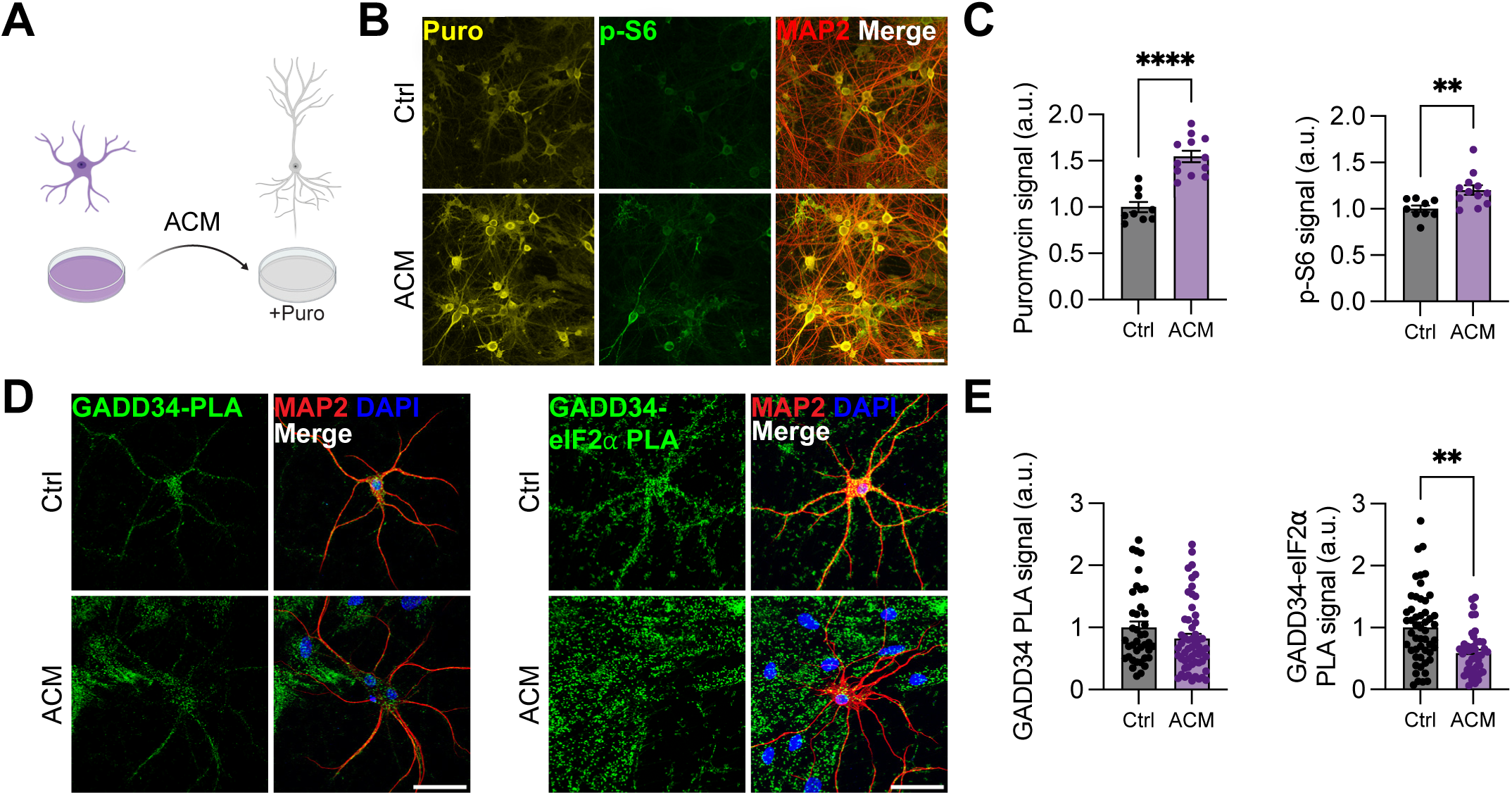
Astrocytes secrete factors that regulate neuronal translation. A. ACM collected from astrocyte cultures was used to treat neural cultures. Puromycin was acutely added to label newly synthesized proteins. B. Representative images of puromycin incorporation and p-S6 signal in MAP2+ control and ACM-treated neurons. Scale bar=100um. C. Quantification of puromycin and p-S6 signal as in panel (B). ACM treatment increases neuronal puromycin and p-S6 signal. Student’s unpaired t test ****p<0.0001, **p<0.01, n=3. Bar plots represent mean ± SEM. D. Representative images of single protein GADD34 PLA and GADD34-eIF2*a* PLA signal in MAP2+ control and ACM-treated neurons. Scale bar=50um. E. Quantification of GADD34 and GADD34-eIF2*a* PLA signal as in panel (D). ACM treatment decreases GADD34-eIF2*a* signal without changing GADD34 signal. Dunnett’s multiple comparisons test, **p<0.01, n=4. Bar plots represent mean ± SEM. Statistics derived from one-way ANOVA across all four conditions, see Fig S15.

To gain insight into the size of astrocyte-secreted factors that regulate increased neuronal translation, we began by fractionating complete ACM using spin columns, generating a “concentrate” and “eluate” that contain molecules larger or smaller than 3kDa, respectively (Fig 2A). Neurons treated with the >3kDa concentrate, but not <3kDa eluate, displayed a similar increase in global translation to that induced by complete ACM (Fig 2B-C). We then sequentially fractionated a new set of ACM through spin columns with 100kDa and 3kDa molecular weight cutoffs (Fig 2D). Neurons treated with either the >100kDa fraction or 3-100kDa fraction display no significant difference in translation compared to non-treated controls (Fig 2E, S3C). These results suggest that there is likely more than one astrocyte-secreted factor that regulates neuronal translation, that the factors vary in size but at minimum are short peptides, and that the effects of these factors are synergistic.

**Figure 2.**
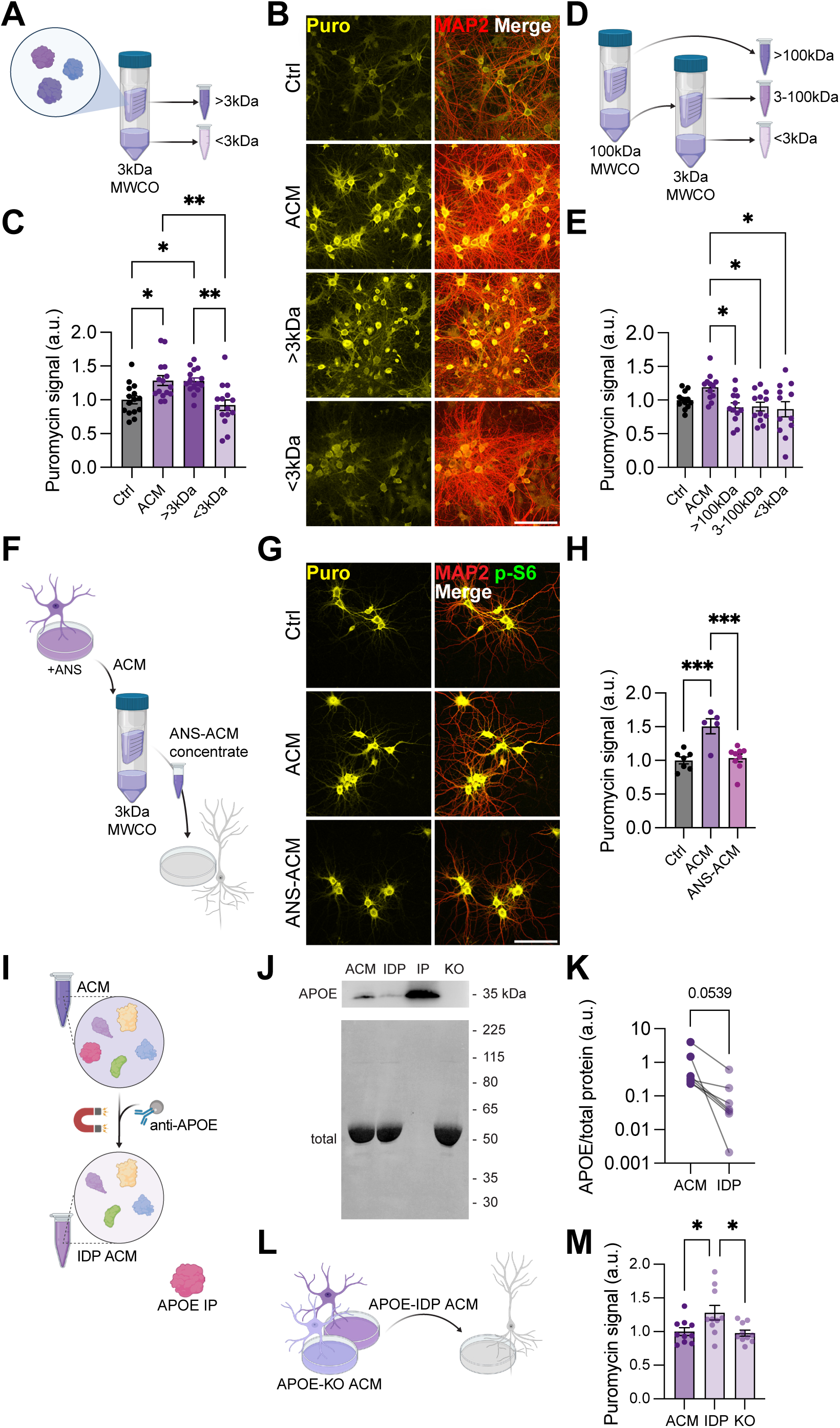
Astrocyte secreted proteins regulate neuronal translation. A. ACM was fractionated through a 3kDa MWCO spin column, then the concentrate and eluate used to treat neurons. B. Representative images of puromycin incorporation in control, ACM, ACM concentrate, or ACM eluate treated neurons. Scale bar=100um. C. Quantification of panel puromycin signal as in panel (B). ACM >3KDa concentrate, but not <3kDa eluate increases neuronal puromycin incorporation. Ordinary one-way ANOVA, Tukey’s multiple comparisons test **p<0.01, *p<0.05, n=5. Bar plots represent mean ± SEM. D. ACM was sequentially fractionated through 100kDa then 3kDa spin columns, then the resulting three fractions were used to treat neurons. E. Quantification of puromycin signal, representative images in Fig S3C. No single fraction is sufficient to increase neuronal puromycin incorporation. Ordinary one-way ANOVA, Tukey’s multiple comparisons test *p<0.05, n=4. Bar plots represent mean ± SEM. Quantification of p-S6 signal in Fig S3D. F. Astrocytes were treated with anisomycin (ANS) prior to collecting ACM, whose concentrate was used to treat neurons. G. Representative images of puromycin incorporation in control, ACM, or ANS-ACM treated neurons. Scale bar=100um. H. Quantification of puromycin signal. ANS-ACM decreases neuronal puromycin incorporation. Ordinary one-way ANOVA, Tukey’s multiple comparisons test, n=3. Bar plots represent mean ± SEM. Quantification of p-S6 signal in Fig S4F. I. Apolipoprotein E (APOE) is an abundant astrocyte-secreted protein. Magnetic beads conjugated with anti-APOE antibodies were used to immunodeplete (IDP) APOE from ACM. J. Representative western blot of APOE before and after IDP. APOE content is decreased in the IDP fraction, but present in the immunoprecitate (IP). No APOE is found in ACM generated from APOE knockout (KO) astrocytes. K. Quantification of APOE signal as in panel (I). APOE content in ACM decreases after immunodepletion. Student’s paired t test, n=7. Bar plots represent mean ± SEM. L. APOE-IDP or APOE-KO ACM was used to treat neurons. M. Quantification of puromycin signal. APOE-IDP, but not KO ACM, further increases neuronal translation as compared to control ACM. Ordinary one-way ANOVA, Tukey’s multiple comparisons test *p<0.05, n=3. Bar plots represent mean ± SEM.

Given that proteins are significantly enriched in spin column concentrate (Fig S3A), we next asked whether the ACM factors that regulate neuronal translation are proteinaceous. We used low-dose anisomycin to inhibit protein synthesis for 24h prior to ACM collection (Fig 2F). Filtering this anisomycin-ACM (ANS-ACM) through spin columns removed any excess anisomycin and prevented its direct effects on neurons (Fig S4A-C). Neurons treated with this ANS-ACM concentrate displayed no significant difference in translation compared to non-treated controls (Fig 2G-H, S4D-F). These results suggest that the factors mediating this astrocyte-neuron regulation include newly synthesized astrocytic proteins, and given the results of size-based fractionation, these proteins are potentially bound to larger complexes such as lipoproteins or extracellular vesicles.

Apolipoprotein E (APOE) is one of the most abundant proteins secreted by astrocytes^48^. We reasoned that APOE may be a good candidate to further examine because it is known to stabilize lipoprotein structures and transport its cargo in between cells^49^. Furthermore, although free APOE is around 36kDa, a lipoprotein can be very large, matching both the 3-100kDa and >100kDa size ranges we previously established. Indeed, we detected APOE in our >3kDa ACM concentrate (Fig S5A). We began by examining how neuronal translation would change by removing APOE from ACM. Using anti-APOE antibodies, endogenous APOE was largely bound to magnetic beads (immunoprecipitated, IP fraction) and immunodepleted from ACM (IDP fraction) (Fig 2I-K). This IDP-ACM led to a further increase in neuronal translation compared to complete ACM (Fig 2L-M), suggesting that the fraction bound to APOE usually has an inhibitory effect. On the other hand, neurons treated with ACM generated from APOE knockout mice (APOE KO) did not differ in translation from those treated with complete ACM (Fig 2M). Given the constitutive nature of this knockout, however, we cannot rule out compensatory upregulation of other apolipoproteins or lipid carriers across development, and we therefore interpret this result cautiously. To more directly test whether APOE protein itself drives translational inhibition, we treated neurons with recombinant APOE (rAPOE) purchased from two separate vendors. Dosage was determined by measuring endogenous APOE concentrations from ACM samples via ELISA (Fig S5C). Surprisingly, rAPOE led to similar increases in neuronal translation compared to ACM-treated neurons (Fig S5D, center), and this increase was dose-dependent (Fig S5D, right). This suggests that APOE in its free and bound forms may have different functions, consistent with previous reports^50,51,49^. To determine which function may be more relevant for our observations, we next asked whether the APOE found in our ACM is free or bound. APOE immunoprecipitated from ACM runs at a larger than expected size (∼100kDa) without additional detergent, suggesting that this APOE is packaged within a larger complex (Fig S5E). To confirm that free APOE is not the relevant form in our model, we took advantage of the fact that LDL receptor related protein associated protein 1 (LRPAP1) is a competitive inhibitor of APOE that preferentially binds low-density lipoprotein receptor-related protein 1 (LRP1) and very-low density-lipoprotein receptor (VLDLR)^52^, both of which have high affinity for free but not bound APOE^51^. Specifically, translation in neurons treated with ACM and recombinant LRPAP1 did not differ from those treated with ACM alone (Fig S5F). On the other hand, LRPAP1 effectively blocked increased translation driven by rAPOE, but had no effect on translation alone (Fig S5G). These results indicate that the APOE found in our ACM is primarily packaged in a large complex, consistent with previous reports^53^, and its contents negatively regulate neuronal translation.

### Neurotoxic reactive astrocytes dysregulate neuronal translation

Astrocytes are highly receptive to environmental changes, and in response can enter multiple different types of states and substates^54^. For example, neurotoxic reactivity is a state of inflammation that prevents astrocytes from performing the normal supportive functions usually observed in healthy conditions, and instead these astrocytes are neurotoxic^55^. Importantly, neurotoxic reactive (NR) astrocytes have been observed in multiple neurodegenerative diseases including Alzheimer’s disease^41^. Therefore, we next sought to determine how NR astrocytes impact neuronal translation. Consistent with previous studies^41,55^, the NR astrocytes we generated expressed increased levels of GFAP and C3 and morphologically exhibited increased number and length of branches compared to control counterparts (Fig S6A-C). NR astrocyte conditioned media (NR-ACM) led to decreased neuronal translation compared to control ACM (Fig 3B-C), suggesting abrogation of the previously observed ACM-induced increase. Importantly, direct treatment of neurons with TNF, IL1α, and C1q, the cytokines used to generate NR astrocytes, did not alter translation levels (Fig S6D-E), indicating that the effects observed are due to neurons responding to NR-ACM rather than the cytokines. Consistent with decreased global translation, neurons treated with NR-ACM also displayed decreased p-S6 (Fig 3C). Although total GADD34 levels did not differ between neurons treated with ACM and NR-ACM, the GADD34-eIF2α interaction was significantly increased in neurons treated with NR-ACM (Fig 3D-E, S7). Given that NR astrocytes have been previously reported to overexpress and oversecrete APOE^53^, we asked whether targeting APOE can rescue the decrease in translation. Neurons treated with APOE-immunodepleted NR-ACM (IDP-NR-ACM) displayed no significant difference in translation as compared to those treated with control ACM (Fig 3H-I), indicating effective rescue of the toxic effect.

**Figure 3.**
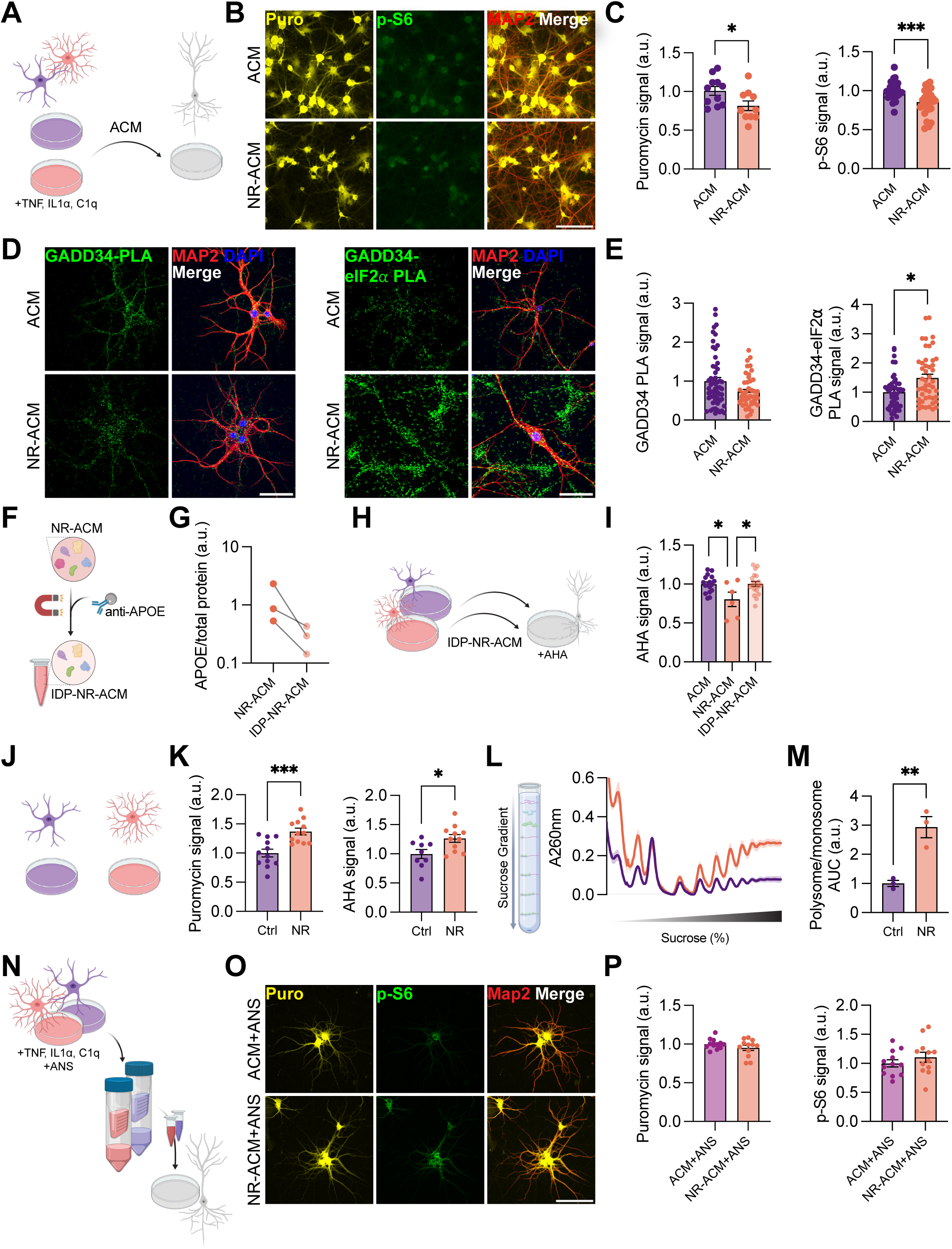
Neurotoxic reactive astrocytes dysregulate neuronal translation. A. TNF, IL1*a*, and C1q were used to generate neurotoxic reactive (NR) astrocytes. Control ACM or NR-ACM was then used to treat neurons. B. Representative images of puromycin incorporation and p-S6 signal in control ACM and NR-ACM treated neurons. Scale bar=100um. C. Quantification of puromycin and p-S6 signal as in panel (B). NR-ACM decreases neuronal puromycin and p-S6 signal as compared to control ACM. Student’s unpaired t test, ***p<0.001, *p<0.05, puromycin n=3, p-S6 n=9. Bar plots represent mean ± SEM. D. Representative images of single protein GADD34 and GADD34-eIF2*a* PLA signal in MAP2+ control ACM and NR-ACM treated neurons. E. Quantification of GADD34 and GADD34-eIF2*a* PLA signal as in panel (D). NR-ACM treatment increases GADD34-eIF2*a* without changing GADD34 signal. Dunnett’s multiple comparisons test, *p<0.05, n=4. Bar plots represent mean ± SEM. Statistics derived from one-way ANOVA across all four conditions, see Fig S15. F. Magnetic beads conjugated with anti-APOE antibodies were used to immunodeplete (IDP) APOE from NR-ACM, producing APOE-IDP NR-ACM (IDP-NR-ACM). G. Quantification of APOE signal. APOE content in NR-ACM decreases after immunodepletion. Bar plots represent mean ± SEM. H. Control ACM, complete NR-ACM, or IDP-NR-ACM was used to treat neurons. L-azidohomoalanine (AHA) was acutely added to label newly synthesized proteins. I. Quantification of AHA signal. IDP-NR-ACM induces no difference in neuronal AHA incorporation as compared to control ACM. Ordinary one-way ANOVA, Tukey’s multiple comparisons test *p<0.05, ACM and IDP-NR-ACM n=6, NR-ACM n=2. Bar plots represent mean ± SEM. J. TNF, IL1*a*, and C1q were used to generate neurotoxic reactive (NR) astrocytes. K. Quantification of puromycin signal and AHA signal in control or NR astrocytes. NR astrocytes display increased puromycin and AHA incorporation as compared to control astrocytes. Student’s unpaired t test ***p<0.001, *p<0.05, n=3. Bar plots represent mean ± SEM. Representative images in Fig S8. L. Polysome profiling of control and NR astrocyte ribosome-mRNA complexes, separated by weight on sucrose gradients. Line graph represents mean ± SEM. M. Quantification of area under the curve (AUC) corresponding to the polysome region in panel (M). NR astrocytes display increased polysomes as compared to control astrocytes. Student’s unpaired t test **p<0.01, n=3. Bar plots represent mean ± SEM. N. Astrocytes were treated with anisomycin (ANS) concurrent with NR induction prior to collecting ACM, whose concentrate was used to treat neurons. O. Representative images of puromycin incorporation and p-S6 signal in ACM+ANS or NR-ACM+ANS treated neurons. Scale bar=100um. P. Quantification of puromycin and p-S6 signal in neurons. NR-ACM+ANS induces no difference in neuronal puromycin or p-S6 as compared to control ACM+ANS. Student’s unpaired t test, n=4. Bar plots represent mean ± SEM.

We then turned our attention to the astrocytes themselves. Compared to controls, NR astrocytes displayed increased global translation assayed by puromycin and AHA, as well as increased p-S6 (Fig 3K, S8). To verify this increase, we used polysome profiling, an assay that measures the relative abundance of ribosome-mRNA complexes separated by size on a sucrose gradient^56^. Consistent with our puromycin and AHA incorporation data, NR astrocytes exhibit significantly higher polysome/monosome ratio (Fig 3L-M), indicating higher levels of translation. Given the drastic increase in translation observed, we hypothesized that targeting translation may also be able to rescue the toxic effect of NRAs on neuronal translation. Because ACM enhances neuronal translation in a manner dependent on ongoing astrocyte protein synthesis (Fig 2G-H), we used ACM+ANS as the baseline-equivalent reference. ACM was collected from NR astrocytes treated with anisomycin concurrent with NR induction (NR-ACM+ANS) and concentrated as previously described (Fig 3N). Neurons treated with NR-ACM+ANS concentrate did not differ from ACM+ANS (Fig 3O-P), indicating that the NR-specific suppressive effect on neuronal translation is eliminated when protein synthesis is blocked in NR astrocytes. These results suggest that increased translation is not only a marked phenotype of NR astrocytes but also is required for the suppression of neuronal translation by NR-ACM.

### BDNF stimulated astrocytes differentially regulate neuronal translation

In addition to sensing chronic environmental signals such as inflammation, astrocytes are also sensitive to the rapid kinetics of neuronal activity. To examine how astrocytes might modulate neuronal translation in response to activity, we decided to use brain-derived neurotrophic factor (BDNF), a neurotrophin that is expressed and secreted in response to neuronal activity^57^, and known to induce protein synthesis-dependent forms of plasticity^58^. BDNF has been previously used to model activity and plasticity in vitro^59^, and importantly, astrocytes are known to express the BDNF receptor TrkB^60^. Therefore, we acutely stimulated astrocytes with BDNF before collecting ACM (BDNF-ACM). To eliminate the direct effects of leftover exogenous BDNF in ACM, before treating neurons we also added TrkB-Fc (Fig 4A), a recombinant protein that contains the extracellular domain of TrkB receptor which acts as a scavenger for free BDNF^18^. Compared to its non-targeting control IgG-Fc, TrkB-Fc effectively inhibited direct BDNF activity on neuronal translation (Fig S9A). Neurons treated with BDNF-ACM displayed increased translation and increased p-S6 compared to control ACM (Fig 4B, C). At the same time, BDNF-ACM treated neurons displayed increased GADD34-eIF2α interaction without changing GADD34 levels (Fig 4D-E, S9B-C). We next asked whether the increase in neuronal translation is due to increased synthesis and secretion of astrocyte factors overall. Surprisingly, both our puromycin incorporation and polysome profiling data indicated that astrocytes stimulated with BDNF do not differ in global translation from control astrocytes (Fig 4G-I). Furthermore, total protein content in BDNF-ACM was not different from that of control ACM (Fig 4J). Taken together, these results suggest that without increasing total translational output, astrocytes may be altering their profiles of secreted factors to further increase neuronal translation in response to BDNF activity.

**Figure 4.**
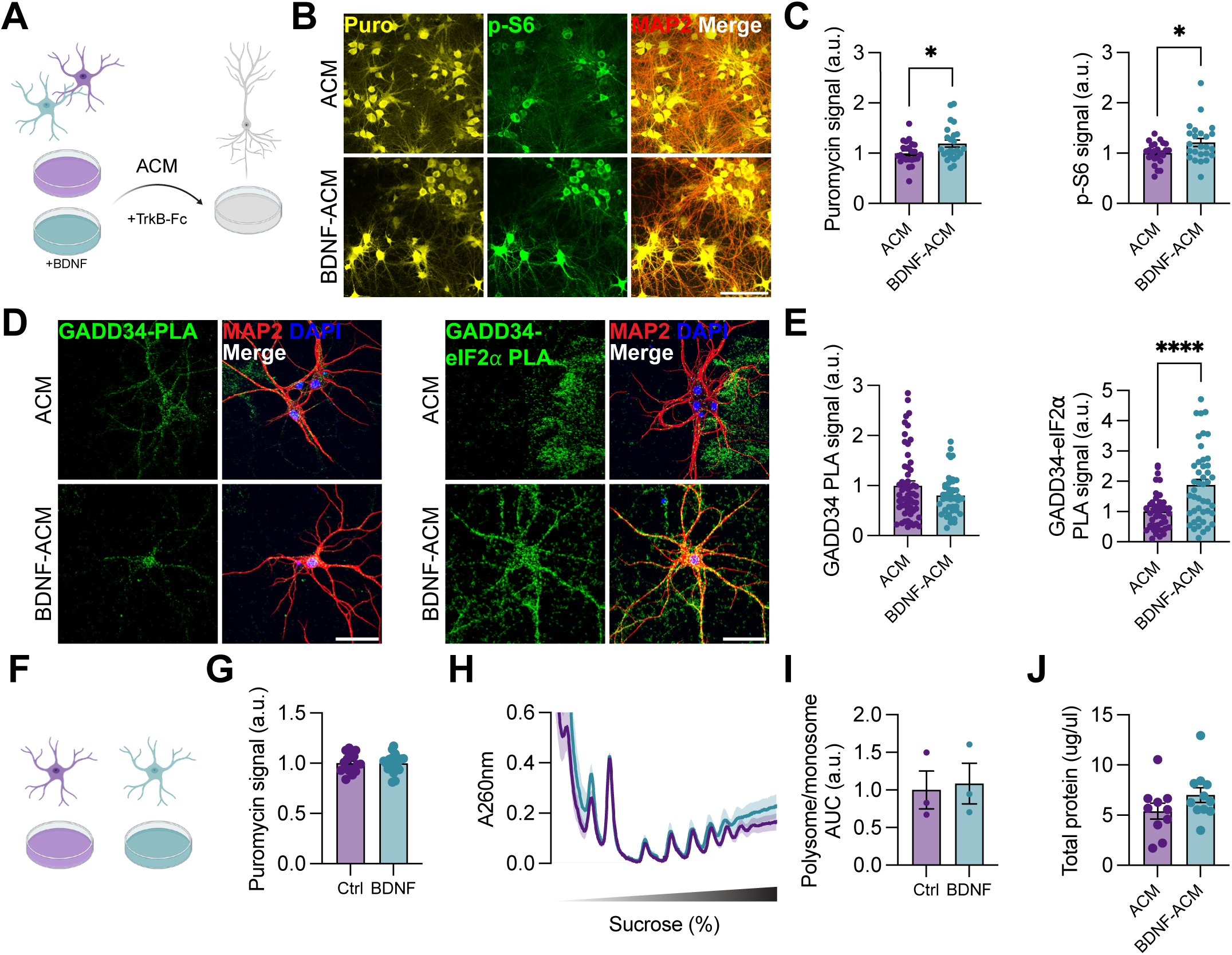
BDNF stimulated astrocytes differentially regulate neuronal translation. A. ACM collected from BDNF-stimulated astrocytes were used to treat neurons. TrkB-Fc was concurrently added to eliminate direct effects of exogenous BDNF. B. Representative images of puromycin incorporation and p-S6 signal in MAP2+ neurons treated with control ACM and BDNF-ACM. Scale bar =100um. C. Quantification of panel (B). BDNF-ACM increases neuronal puromycin and p-S6 as compared to control ACM. Student’s unpaired t test, *p<0.05, n=5. Bar plots represent mean ± SEM. D. Representative images of single protein GADD34 and GADD34-eIF2*a* PLA signal in neurons treated with control ACM and BDNF-ACM. E. Quantification of GADD34 and GADD34-eIF2*a* PLA signal as in panel (D). BDNF-ACM treatment does not alter neuronal GADD34 but increases GADD34-eIF2*a* as compared to control ACM treatment. Dunnett’s multiple comparisons test, ****p<0.0001, n=4. Bar plots represent mean ± SEM. Statistics derived from one-way ANOVA across all four conditions, see Fig S15. F. Astrocytes were stimulated with BDNF, then treated with puromycin to label newly synthesized proteins. G. Quantification of puromycin incorporation in astrocytes. There is no difference in puromycin signal in BDNF-stimulated vs control astrocytes. Student’s unpaired t test, n=6. Bar plots represent mean ± SEM. H. Polysome profiling of ribosome-mRNA complexes from control and BDNF-stimulated astrocytes. I. Quantification of area under the curve (AUC) corresponding to the polysome region in panel (H). There is no difference in the polysome peaks between control and BDNF-stimulated astrocytes. Student’s paired t test, n=3. Line graph represents mean ± SEM. J. Quantification of total protein content in control and BDNF-ACM. There is no difference in protein concentrations between the two groups. Student’s unpaired t test, ACM n=10, BDNF-ACM n=11. Bar plots represent mean ± SEM.

To determine how astrocyte regulation of neuronal translation may be functionally relevant, we turned to morphological analysis of synapses and neuronal structure. Neurons treated with ACM and BDNF-ACM displayed significantly decreased PSD95 and synapsin (SYN1), and their colocalized puncta (Fig 5A-C). At the same time, mean PSD95 and SYN1 intensity also decreased after ACM and BDNF-ACM treatment (Fig S10B). This result was surprising, given that previous literature has reported that ACM promotes synaptogenesis^24,26,61,62^. We first ensured that ACM treatment of our mixed neuron cultures did not lead to higher levels of cleaved-caspase 3, eliminating the possibility of ACM inducing increased cell death (Fig S10C). Furthermore, we determined that our treated neurons displayed no significant changes in any gross morphological characteristics (Fig S10D). We posited that the main difference lay in the fact that in these previous reports pure neuron cultures were used, whereas our cultures already contain many glial cells and are therefore more robust and potentially developmentally more mature. Indeed, when we generated pure neuron cultures, individual and colocalized PSD95/SYN1 puncta increased with ACM treatment as compared to no-ACM controls, replicating previous reports (Fig S10E). These results suggest that in the context of full glial support throughout neuronal development, additional ACM may decrease synapse number to increase synapse specificity.

**Figure 5.**
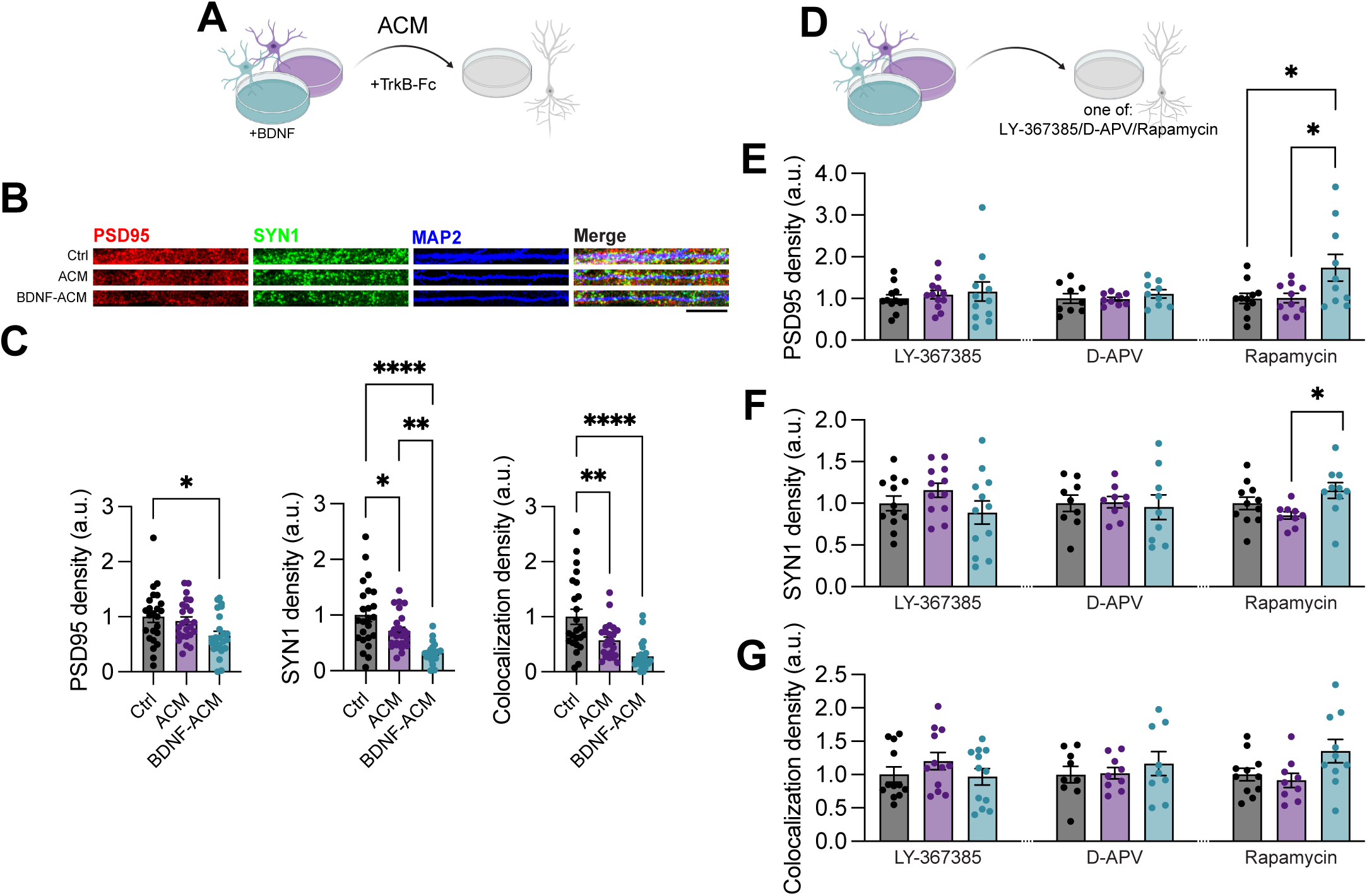
Astrocyte regulation of neuronal translation modulates synapses. A. ACM collected from BDNF-stimulated astrocytes were used to treat neurons. TrkB-Fc was concurrently added to eliminate direct effects of exogenous BDNF. B. Representative images of PSD95 and synapsin (SYN1) signal in neurons treated with control ACM and BDNF-ACM. Scale bar=20um C. Quantification of PSD95 and SYN1 puncta as in panel (B). ACM and BDNF-ACM decrease both individual and colocalized puncta as compared to non-treated controls. Ordinary one-way ANOVA, Tukey’s multiple comparisons test ****p<0.0001, **p<0.01, *p<0.05, n=5. Bar plots represent mean ± SEM. D. Control ACM, BDNF-ACM, and TrkB-Fc were used to treat neurons as in panel (A). Neurons were either concurrently treated with one of LY-367385, D-APV, or rapamycin. Control neurons with no ACM treatment also received one of the above drugs. E. Quantification of individual PSD95 puncta in neurons. LY-367385 and D-APV block the ACM and BDNF-ACM driven decrease in PSD95, and rapamycin further leads to an increase in PSD95 in BDNF-ACM treated neurons. Separate ordinary one-way ANOVAs performed for each drug/shRNA treatment, Tukey’s multiple comparisons test *p<0.05; LY-367385 n=4, D-APV n=3, rapamycin n=4. Bar plots represent mean ± SEM. F. Quantification of individual SYN1 puncta in neurons. LY-367385, D-APV, and rapamycin all block the ACM and BDNF-ACM driven decrease in SYN1. Separate ordinary one-way ANOVAs performed for each treatment, Tukey’s multiple comparisons test *p<0.05; LY-367385 n=4, D-APV n=3, rapamycin n=4. Bar plots represent mean ± SEM. G. Quantification of colocalized PSD95/SYN1 puncta in neurons. LY-367385, D-APV, and rapamycin all block the ACM and BDNF-ACM driven decrease in colocalized puncta. Separate ordinary one-way ANOVAs performed for each treatment, LY-367385 n=4, D-APV n=3, rapamycin n=4. Bar plots represent mean ± SEM.

One explanation for the observed decrease in synapses is synaptic pruning. To examine this possibility, we collected total RNA samples from our mixed neural cells with or without ACM treatment, as well as from astrocytes with or without BDNF treatment, to measure relative abundance of specific mRNA transcripts related to pruning via quantitative real-time PCR (qRT-PCR). We examined opsonization signals such as *C3* and *C1q*, which promote pruning by microglia^63^, as well as astrocyte phagocytic receptors such as *Megf10* and *Mertk*^27^. Although there were some interesting differences between treatment groups, there was no consistent pattern in the expression of these transcripts to suggest increased synaptic pruning (Fig S11A). Furthermore, although microglia depletion using the CSF1R inhibitor PLX-3397 prevented the ACM-driven decrease in PSD95, it did not block the decrease in synapsin (Fig S11B). Collectively, these data suggest that synaptic pruning is not the likely explanation for the ACM-driven decrease in synapses.

Because glia-mediated synaptic pruning did not account for the ACM-induced reductions in synapses, we next considered whether these changes might instead arise from synapse-intrinsic plasticity mechanisms. Specifically, we hypothesized that our observations involve activity-dependent synaptic weakening mechanisms associated with synaptic depression, which require glutamate receptor signaling and, in some cases, protein synthesis^64^. To test whether glutamate receptor activity is required, we co-treated neurons with ACM and either the mGluR1 antagonist LY-367385^65^ or the NMDAR antagonist D-APV^66^ (Fig 5D). Both inhibitors prevented the ACM-induced loss of synapses (Fig. 5E-G, left two columns). We next asked whether translational control contributes to this process. Across our experiments, we consistently observed translation and p-S6 readouts changing in parallel, pointing to mTORC1 activity as a recurring node for ACM-dependent translational control. Rapamycin not only prevented the ACM-induced decrease in synapses but also led to a net increase in PSD95 and synapsin levels above baseline (Fig. 5E-G, right column), indicating that mTORC1 signaling actively restrains synapse number during this process. Importantly, none of these pharmacological manipulations altered synapses under baseline conditions (Fig S12), indicating that the observed rescues are specific to ACM treatment. Collectively, these findings suggest that ACM induces a coordinated structural refinement of synapses, driven by activity-dependent, mTORC1-regulated translational mechanisms intrinsic to neurons.

Given that astrocyte-secreted APOE affects neuronal translation in physiological and inflammatory contexts (Fig 2N, 2O, 3G), we asked whether APOE is differentially secreted in response to BDNF, and whether this contributes to the observed ACM-induced synaptic remodeling. Compared to control ACM, we found that BDNF-ACM contains significantly higher APOE content (Fig S13A). Removing APOE from ACM and BDNF-ACM via immunodepletion prevented the decrease in synapses that was previously observed (Fig S13B-C). These findings position APOE as a key astrocyte-secreted signal linking translational regulation to synaptic remodeling, though how this function relates to APOE’s role in suppressing global translation remains unresolved. Together, these data implicate astrocyte-secreted APOE as a key signal driving the activity-dependent, translation-regulated synaptic remodeling described above, linking a specific astrocyte-derived factor to a defined neuronal mechanism of structural plasticity.

### Endocytosis mediates ACM effects on neuronal translation

Given the robust effects of astrocyte secreted proteins in affecting neuronal translation, we next asked how these extracellular signals are relayed to neurons. One hypothesis is that the secreted factors are endocytosed and trafficked directly into neurons. To examine this possibility, we treated neurons with control ACM, BDNF-ACM, or NR-ACM, in conjunction with Dynasore, a dynamin inhibitor that blocks endocytic vesicle pinching^67^ (Fig 6A). Dynasore effectively blocked not only ACM and BDNF-ACM mediated increases in translation, but also blocked the decrease mediated by NR-ACM (Fig 6B-C). At the same time, Dynasore effectively blocked ACM and BDNF-ACM mediated decreases in synapses (Fig 6D-E). Neurotoxic reactive astrocytes have been previously reported to reduce synapse number^41^: we found that Dynasore treatment also rescued NR-ACM mediated decrease in synapses (Fig 6E). Treatment using a different endocytosis inhibitor, Pitstop2^68^, similarly blocked ACM induced changes in neuronal translation and synapses (Fig S14A-C). Dynasore and Pitstop2 do not affect global translation or synapses at baseline (Fig S14D). These results indicate that endocytic uptake of astrocyte-secreted cargo is required for neurons to relay both translation-promoting and translation-suppressing signals across distinct astrocyte states, pointing to a shared internalization step as a convergent node through which diverse extracellular cues reach the neuronal translational machinery. Collectively, these results demonstrate that astrocyte-secreted factors act through neuronal endocytosis and mTORC1/ISR signaling to differentially regulate neuronal translation and synaptic structure across astrocyte states.

**Figure 6.**
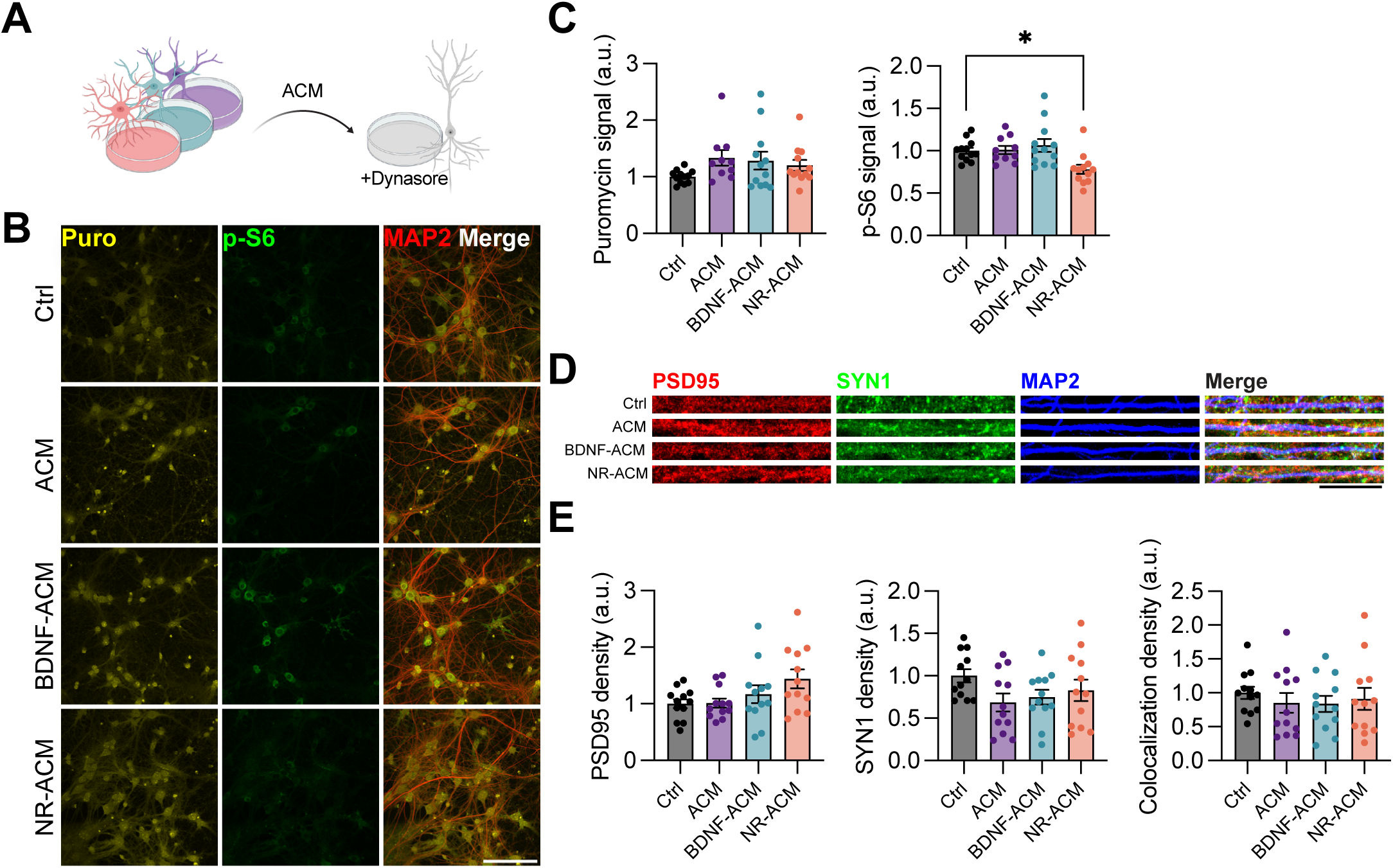
Endocytosis mediates ACM effects on neuronal translation. A. Control ACM, BDNF-ACM, and NR-ACM were used to treat neurons. Neurons were concurrently treated with Dynasore. B. Representative images of puromycin incorporation and p-S6 signal in neurons treated with Dynasore and different types of ACM. Control neurons with no ACM treatment also received Dynasore. Scale bar=100um. C. Quantification of puromycin and p-S6 signal in neurons as in panel (B). Dynasore blocks increases in neuronal puromycin and p-S6 driven by control ACM and BDNF-ACM, while also blocking the decrease in neuronal puromycin and p-S6 driven by NR-ACM. Ordinary one-way ANOVA, Tukey’s multiple comparisons test, *p<0.05, n=4. Bar plots represent mean ± SEM. D. Representative images of PSD95 and SYN1 signal in neurons treated with Dynasore and different types of ACM. Control neurons with no ACM treatment also received Dynasore. Scale bar=20um. E. Quantification of PSD95 and SYN1 puncta as in panel (D). Dynasore blocks decreases in individual and colocalized PSD95/SYN1 puncta driven by control ACM, BDNF-ACM, and NR-ACM. Ordinary one-way ANOVA, n=4. Bar plots represent mean ± SEM.

## Discussion

Herein, we identify astrocyte-secreted proteins as regulators of neuronal translation and demonstrate that astrocytes can differentially control neuronal protein synthesis across physiological, activity-dependent, and disease-associated states. Although neuronal protein synthesis has long been known to be critical for the formation of long-lasting synaptic plasticity and long-term memory, whether its regulation is non-cell autonomous or otherwise is poorly understood. Our findings show that astrocyte-derived signals influence not only neuronal translational output but also neuronal mTORC1 and ISR signaling, placing astrocytes as active regulators of neuronal proteostatic pathways.

Several observations support an instructive role for this novel astrocyte-to-neuron signaling. Across distinct astrocyte states, neuronal translational responses are not predicted by global astrocyte translational activity. In inflammation, neurotoxic reactive astrocytes markedly suppressed neuronal translation, despite increased astrocyte translation, confirmed by both puromycin/AHA incorporation and polysome profiling. On the other hand, BDNF-stimulated astrocytes enhanced neuronal translation despite no detectable change in global astrocyte translation. If astrocytes simply provided permissive, generic translational support to neurons, neuronal translational output might be expected to scale with astrocytes’ own translational and secretory capacity; instead, the direction and magnitude of the neuronal response varied independently of astrocyte translation, suggesting that the content, rather than the quantity, of astrocyte-secreted signals determines the neuronal outcome. Separately, we identified an astrocyte-secreted negative regulator of neuronal translation, APOE, that acts across astrocyte states. Together, this lack of correlation between astrocyte and neuronal translational responses, combined with the identified existence of negative regulators, is difficult to reconcile with a purely permissive model and instead suggests that astrocytes actively instruct neuronal translational programs.

The signaling pathways underlying these effects further support a model of state-specific, instructive control rather than a single uniform translational switch. mTORC1 activity, assessed by p-S6, tracked consistently with translational output across every astrocyte state we examined – increasing with ACM and BDNF-ACM, and decreasing with NR-ACM – positioning mTORC1 as a core node of convergence through which diverse astrocyte-derived signals tune neuronal translation. In contrast, GADD34 activity did not follow a single predictable relationship with translational output. Relative to control, ACM decreased GADD34-eIF2α interactions despite increasing translation, opposite to the relationship expected if GADD34 relieves p-eIF2α-mediated translational repression. Moreover, relative to ACM, both BDNF-ACM and NR-ACM significantly increased GADD34-eIF2α interactions yet had opposite effects on translation: BDNF-ACM caused increases and NR-ACM caused decreases. Notably, total GADD34 levels did not significantly differ across any of these conditions, indicating that these changes in interaction are not simply a function of GADD34 abundance. These results show that the same direction of change in GADD34-eIF2α interactions can accompany opposite translational outcomes, suggesting that astrocyte-derived signals likely engage the ISR through distinct, state-specific mechanisms rather than a single shared biochemical logic. At the same time, our results indicate that GADD34-eIF2α interaction is not, on its own, a reliable predictor of net translational output. Directly assaying corresponding changes in eIF2α phosphorylation status, the canonical functional readout of ISR activity, will be necessary to clarify this relationship. Despite this limitation, these findings suggest that astrocytes instructively shape neuronal translation by coordinating a convergent mTORC1 signal with state-specific engagement of the ISR, allowing distinct astrocyte-derived cues to produce tailored translational outcomes in neurons.

One way that astrocytes implement this instructive control is through APOE and its associated cargo. In basal conditions, we found that depletion of APOE and its cargo causes an increase in translation relative to ACM-induced levels, suggesting that when present, these factors induce a negative regulatory effect. Consistent with this inference, neurotoxic reactive astrocytes are known to increase APOE secretion, and we observed that depletion can rescue the decreased neuronal translation induced by neurotoxic reactive ACM. These findings define a specific astrocyte-derived signaling mechanism that contributes to the negative regulation of neuronal protein synthesis. Notably, we found that the negative effect is not due to free APOE alone, instead implicating its associated cargo; indeed, recombinant free APOE alone increased rather than decreased neuronal translation, opposite to the inhibitory effect of ACM with predominantly bound, complexed APOE. Further studies are required to dissect exactly which factors are involved: a candidate could be screened based on the recently published lipidome and proteome of astrocyte-secreted APOE-bound lipoproteins^69^.

More broadly, this novel role of APOE has important implications for disease. APOE is one of the most widely studied genes in cardiovascular health and neurodegenerative disorders, primarily because the human APOE4 variant is associated with higher genetic risk for atherosclerosis and Alzheimer’s disease^70^. In animal models, mice carrying human APOE4 have been shown to display impaired memory that can be rescued by injecting drugs that target the ISR to generally increase translation^71^, however the cell type-specific effects of APOE4 on translation are poorly understood. Future studies could leverage isogenic human induced pluripotent stem cells with different APOE variants^72^ to develop neuron and astrocyte models with mixed APOE background to examine how neuronal translation, and its regulation by astrocytes, changes solely due to genotype.

The effects of APOE on neuronal translation are further shaped by its receptors, which belong to the low-density lipoprotein receptor (LDLR) family and bind APOE with affinity that varies by variant and lipidation status^51^. This receptor diversity could help explain why free and bound APOE produce opposite effects on neuronal translation in our system. For example, APOE4 has been shown to impair LRP8 (APOER2) recycling, disrupting glutamate receptor trafficking and Reelin-dependent LTP^73^, a mechanism that, if relevant here, could link APOE receptor biology directly to the synaptic phenotypes we observed. Altering the expression pattern of different APOE receptors may therefore provide an additional means of fine-tuning the neuronal response to astrocyte-secreted APOE.

Mechanistically, we found that astrocyte-secreted proteins require neuronal endocytosis to influence translational control. This observation is consistent with the known role of APOE serving as a transport molecule specifically through receptor-mediated endocytosis^73^. Furthermore, it suggests that rather than remaining at the cell membrane, astrocytic proteins could traffic into neurons to participate directly in neuronal mTORC1 or ISR signaling. Indeed, many studies have previously identified multiple species of translation factors, ribosomal proteins, or translation regulation elements that are present in ACM^48,74^, and the abundance of these factors can change in response to different forms of stimulation or astrocyte states^22,53,75^. Whether or not these astrocyte-secreted translational regulators truly enter neurons remains unknown. It is also unknown whether these factors are secreted as free proteins, or potentially in the form of extracellular vesicles or APOE-bound lipoproteins. Regardless of the precise route, this requirement for endocytosis links astrocyte-derived signals to downstream translational and structural changes in neurons.

Functionally, we found that the effects on neuronal translation caused by astrocytes are accompanied by changes in synapse number that depend on glutamatergic activity as well as neuronal mTORC1 signaling. These requirements are in line with known activity-dependent synaptic weakening mechanisms associated with synaptic depression, suggesting that astrocyte-secreted proteins may help to refine synaptic networks. These data also provide a novel translation-based mechanism for activity-driven synaptic remodeling by astrocytes^76^.

Within this mechanism, however, APOE presents a notable exception. Although APOE is involved in negatively regulating global neuronal translation, its depletion also prevented ACM-induced synapse loss, presenting an unexpected puzzle where a single factor is linked to opposing roles in global translational output versus synaptic remodeling. Notably, APOE receptors localize prominently to synapses^77^, at both the presynaptic membrane and postsynaptic density^78^, suggesting that APOE signaling is concentrated at synaptic sites in a manner not captured by our bulk measurements. Indeed, local translation at dendrites and synapses is known to play a critical role in synaptic plasticity^58,79^, raising the possibility that APOE preferentially acts on this local translational machinery rather than, or in addition to, global translation. Alternatively, APOE-amyloid-β complexes have been shown to accumulate in synapses, leading to synapse loss in AD^80^, pointing to a translation-independent mechanism in which APOE-bound cargo accumulates directly at the synapse to drive synapse loss. Examining these potential mechanisms, particularly through subcellular fractionation or compartment-restricted labeling approaches, will be an important direction for future work.

Our findings establish a conceptual framework for how astrocytes actively, rather than passively, shape neuronal proteostasis, and raise several broader questions for future investigation. Having identified APOE as a negative regulator of neuronal translation, an open question is the identity of positive astrocyte-secreted factors and whether their balance with negative factors shifts across astrocyte states. Unbiased proteomic profiling of secreted and neuronally endocytosed proteins will help define this full repertoire of signals. In addition, our bulk translation readouts cannot resolve which specific transcripts are regulated. Sequencing the neuronal translatome could reveal whether astrocytes selectively tune translation of specific transcript classes, such as synaptic or plasticity-related mRNAs, and whether mTORC1- and ISR-dependent effects act on shared or distinct targets. We also focused exclusively on astrocyte-secreted, diffusible signals; future studies incorporating controlled astrocyte-neuron co-culture or in vivo manipulation will be important for determining how adhesion molecule signaling, neurotransmitter recycling, ion buffering, and astrocyte heterogeneity across circuits and brain regions contribute to or interact with the secreted signals described here. Finally, our findings raise the possibility that targeting astrocytic translation or APOE signaling could ameliorate memory impairments seen in AD mouse models.

In conclusion, our findings position astrocytes as active, instructive regulators of neuronal proteostasis, a capacity that may also be redirected to drive pathology in neurodegenerative disease.

## Methods

### Timed Pregnancies

Male and female wild-type or APOE KO (Jackson #000664, #002052) C57BL/6J mice aged 8+ weeks were placed together, then separated 2-3 days after copulation.

### Primary Neuron Preparation

Timed pregnant dams were euthanized and dissected to obtain E17 cortical tissue. Following removal of meninges, tissue was cut into ∼1mm^3^ pieces and digested in 0.25% trypsin-EDTA (Gibco 25200056) for 5min at 37°C, washed thoroughly in DMEM + 1X GlutaMAX + 10% FBS + 1X P/S (DMEM/FBS media) then triturated and passed through a 40um cell strainer. After pelleting and resuspending, cells were plated in DMEM/FBS at 5x10^4^ cells per PLO-coated 12mm coverslip in 24-well plates, or at 1x10^6^ cells per PLO-coated 10cm dish. One to two hours after plating, once cells had settled, media was switched to Neurobasal Plus + B27 Plus + 0.25X GlutaMAX + 1X P/S (NB/B27+ media). Cultures were fed twice a week with NB/B27+ and maintained in 37°C CO_2_ incubators for 14 days prior to experiments. For indicated pure neuron cultures, cytosine arabinoside (Ara-C, Sigma C1768) was added to culture medium on DIV3 to a 1μM final concentration, then neurons collected at DIV7. Each neuron prep was considered a separate biological replicate.

### Primary Astrocyte Preparation

Postnatal day 3 (P3) pups were sacrificed to obtain the cortex. Using a tissue dissociation kit (Worthington Biochemical LK003150), the meninges-free, cut tissue pieces were digested with papain for 20min, thoroughly washed in DMEM/FBS, lightly triturated using a 5mL serological pipette in ovomucoid inhibitor, passed through a 40μm cell strainer, then pelleted and resuspended with a solution of 0.5% IgG-free BSA (Jackson ImmunoResearch 001-000-161) in PBS. Adapting protocols from Holt & Olsen 2016^81^ and Miltenyi Biotec MACS Cell Separation kits, microglia and myelin debris were concurrently depleted using anti-CD11b and anti-myelin microbeads, then astrocytes were selected using anti-ACSA-2 microbeads (Miltenyi Biotec 130-093-634, 130-096-733, 130-097-678). Purified astrocytes were plated on PLO-coated 10cm dishes at 1x10^6^cells/dish, and cultured in NB/B27+ supplemented with 5ng/μL HB-EGF^82^ (PeproTech 100-47). Cultures were fed twice a week and maintained for at least one week prior to experiments. Each astrocyte prep was considered a biological replicate.

### ACM Generation

All forms of ACM were collected after 72h of conditioning, i.e. three days after fresh media exchange. For NR-ACM, 3ng/mL Il-1α (PeproTech AF-200-01A) + 30ng/mL TNF (PeproTech 300-01A) + 400ng/mL C1q (Complement Technology A099) was spiked into cultures 24h prior to ACM collection. For BDNF-ACM, 50ng/mL BDNF (StemCell Technologies 78005.1) was spiked into cultures 1h prior to ACM collection. For ANS-ACM, 1uM anisomycin (Sigma A9789) was spiked into cultures 24h prior to ACM collection. Collected ACM was stored frozen at -20°C until use in experiments.

### ACM Concentration

ACM was centrifuged at ∼3,000g for 2h at 4°C through 3kDa or 100kDa MWCO spin columns (Cytiva 28932358, 28932363). The resulting concentrate is less than 10% of the original ACM volume, and both concentrate and eluate were both collected for use in experiments.

### APOE IDP

ACM was depleted of APOE using the Dynabeads Co-Immunoprecipitation Kit (Invitrogen 14321D) following manufacturers protocol. Briefly, APOE antibody (Novus Biologicals NB110-60531) was coupled to magnetic beads on a rotator at 37°C overnight, washed, then incubated in ACM on a rotator at 4°C overnight. APOE IDP-ACM was then collected by separating beads from supernatant on a magnetic rack. Beads were washed three times in buffer EB (1X IP + 100mM NaCl), once in buffer LWB (1X LWB+0.02% Tween-20), then APOE was eluted in RIPA buffer by incubating for 5min at room temperature followed by 10min at 60°C.

### ACM Treatment

Neuron culture medium was fully removed and replaced with warmed ACM. When indicated, 200ng/mL TrkB-Fc (R&D Systems 688-TK-100) was added to ACM prior to treatment. For controls, old media was pipetted up then down into the same well to physically mimic media exchange. Neurons were treated with ACM for 24h prior to sample collection.

### De novo Protein Synthesis Labeling

Immediately prior to sample collection, cell cultures were spiked with 0.5ug/mL puromycin (Sigma P8833) for 5min, or 4mM or 1mM AHA (Vector Laboratories CCT-1066) for 2h or 24h respectively. Cultures spiked with 10ug/ml cycloheximide for 1h served as labeling controls.

### eGFP Transfection for Morphology

Neurons were sparsely transfected with 100ng of eGFP mRNA (TriLink Biotechnologies L-7601) in 1μL Lipofectamine 2000 (Invitrogen 11668019) and 50μL OptiMEM (Gibco 31985062) per well 4h prior to sample collection.

### APOE ELISA

Endogenous APOE concentration in ACM was measured using the Mouse Apolipoprotein E ELISA Kit (Abcam ab215086) following manufacturers protocol. Briefly, samples were diluted 100X, then standards and samples were incubated with the antibody cocktail on the assay plate for 1h at room temperature. Wells were washed, developed with manufacturers TMB development solution, then stopped and results read on a plate reader at 450nm. Sample concentrations were calculated based on the linear standard curve.

### rAPOE/rLRPAP1 Treatment

Neurons were treated for 24 hours with recombinant mouse APOE (Abcam ab226314, Acro Biosystems APE-M52H5) or recombinant mouse LRPAP1 (Abcam ab201883, R&D Systems 4480-LR) separately or concurrently, at 300ng/mL unless otherwise indicated.

### Pharmacology

Neurons were treated for 24 hours with 10μM LY-367385 (Sigma L4420), 10μM D-APV (Tocris 0106), 20nM rapamycin (Sigma 553210), 10μM Dynasore (Sigma 324410), or 10μM Pitstop2 (Sigma SML1169), then collected at DIV14. For microglia depletion, 1μM PLX-3397 (MedChemExpress HY-16749) was added to culture medium on DIV3 and until sample collection.

### qRT-PCR

Cell lysate was collected in TRIzol Reagent (Invitrogen 15596026), then RNA purified using Direct-zol RNA Miniprep Plus (Zymo Research R2070). Sample cDNA was generated using iScript Reverse Transcription Supermix (Bio-Rad 1708841) following manufacturers protocol and then diluted 10X. Target cDNA were amplified using iTaq Universal SYBR Green Supermix (Bio-Rad 1725120) and the following primers:

GAPDH_FWD: AACTTTGTCAAGCTCATTTCCTGG

GAPDH_REV: TTGGGATAGGGCCTCTCTTGCT

IL33_FWD: CTCACTGCAGGAAAGTACAGC

IL33_REV: GGTCTTCTGTTGGGATCTTCTT

C1qA_FWD: TCCAGTTTGATCGGACCACG

C1qA_REV: GGATTTCCTGGAGCCCCATC

C3_FWD: ATAAAGAGCCAGCGGCTACA

C3_REV: GGGAGTAATGATGGAATACATGGG

H2K1_FWD: TGAGAAGGAGAAACACAGGTGG

H2K1_REV: GTCACCAAGTCCACTCCAGG

CX3CL1_FWD: CGGCATGACGAAATGCGAAA

CX3CL1_REV: TGTGTCGTCTCCAGGACAATG

MEGF10_FWD: GCCCTGTGCAATGAAGTGTG

MEGF10_REV: AACACTGGCAGGATCTGTCG

MERTK_FWD: AGGAAGGTAGGTCCTGGGTG

MERTK_REV: TCTCGGCAGTGCCTAAGGAT

All samples were run in technical duplicates, their C_T_ values averaged, then fold change calculated from ΔΔC_T_ using GAPDH as normalization.

### Immunocytochemistry

Cells in 24-well plates with coverslips were fixed with 4% PFA in PBS for 20min, permeabilized with 0.1% Triton X-100 in PBS for 10min, then blocked with 5% normal goat serum (NGS) in PBS for 1h on a rocker at room temperature. Primary antibodies were diluted 1:1,000 (puromycin, Sigma MABE343; P-S6, Cell Signaling Technology 2211S; Map2 (gp), Synaptic Systems 188004; Map2 (chk), EnCor Biotechnology CPCA-MAP2; cleaved caspase 3, Cell Signaling Technology 9661S; GFP, Invitrogen G10362; GFAP, EMD Millipore AB5541) or 1:500 (PSD95, Synaptic Systems 124308; Syn1, Synaptic Systems 106011) in 5% NGS and incubated on coverslips at 4°C overnight. After PBS washing, coverslips were then incubated with secondary antibodies diluted 1:1,000 (goat anti-guinea pig AF647, goat anti-mouse AF568, goat anti-rabbit AF488, goat anti-chicken AF405; Invitrogen A21450, A11031, A32731, A48260) in 5% NGS for 1h at RT. Coverslips were washed again, then mounted on slides in antifade with or without DAPI (Vector Labs H-1800-10, H-1700-10).

### FUNCAT Click Chemistry

After fixing and permeabilizing (as above), cells on coverslips were blocked with 5% NGS + 5% BSA in PBS for 1h at room temperature. Click reactions were prepared using AF647 alkyne and commercial buffer kits (Vector Labs CCT-1301, CCT-1263) and incubated with samples at 4°C overnight. The next day, coverslips were washed 2X with 0.5mM EDTA in PBS followed by 2X with PBS alone, then blocked and incubated with primary then secondary antibodies as described above.

### PLA Sample Preparation

Proximity Ligation Assay (PLA) was performed using NaveniFlex Cell MR Atto647N (Navinci 60017) following the manufacturer’s suggested protocol. Briefly, cells were fixed and permeabilized as described above, then blocked with Naveni Block solution for 1h at 37°C. The solution was aspirated, replaced with appropriate primary antibody combinations diluted in Naveni Diluent and incubated overnight at 4°C. The following antibody combinations and concentrations were used: rabbit anti-GADD34 (Proteintech 10449-1-AP, 1:500) x mouse anti-eIF2α (Abcam AB5369, 1:300); rabbit anti-GADD34 (Proteintech 10449-1-AP, 1:500) x mouse anti-GADD34 (Abcam ab236516). Guinea-pig anti-MAP2 primary (1:1000) was included alongside all combinations. Coverslips were washed 2x 10 seconds and 1x 15min with TBS-T (0.05% Tween-20) pre-warmed to 37°C. Coverslips were then incubated at 37°C for 60 min in M1 and R2 Navenibodies diluted 1:40 in Naveni Diluent, followed by 2x 10 seconds and 1x 15min washes with pre-warmed TBS-T. Secondary anti-Guinea Pig (1:500) was incubated simultaneously with Navenibodies. Ligation (reaction 1 in standard protocol) was performed at 37°C for 30 min, using standard preparation protocols. Coverslips were washed 1x 10 seconds and then 1x 5min with pre-warmed TBS-T. Rolling circle amplification was performed following standard Navinci reaction preparation directions and incubated at 37°C for 90min. Cells were washed twice with PBS and mounted in antifade with DAPI (Invitrogen P36931).

### Image Acquisition and Analysis

Stained samples were imaged on a Leica SP5 confocal microscope to obtain 8um optical sections with 1um interval thickness. Fiji/ImageJ 2.16.0/1.54p was used for all image processing and quantification using automated scripts. Intensity of signal was normalized against area of MAP2 or GFAP to obtain a mean gray value. PLA puncta in MAP2 signal was quantified using a script developed by Heumüller and colleagues^83^. For analysis of synaptic puncta, Synbot^84^ was used with image segmentation by ilastik, trained on a 5% sample population representing all experimental groups. Total puncta per image were normalized against MAP2 area to obtain puncta density. Two plugins NeurphologyJ^85^ and AnalyzeSkeleton^86^ were used to measure morphology of eGFP-transfected neurons. For all experiments quantifying signal intensity or synaptic puncta, three fields of view were acquired per coverslip, and the average of these three images were plotted as a single data point. For experiments quantifying PLA puncta, five fields of view were acquired per coverslip, and values from each image were plotted as single data points. For experiments quantifying morphological characteristics, values from each individual neuron were plotted as a single data point. Unless otherwise indicated, all values are normalized to the control group mean set to 1 and plotted in arbitrary units (a.u.).

### Western Blotting

Cells were lysed in RIPA buffer (Thermo Scientific 89900) then spun to remove DNA and large debris. Protein concentration was determined by BCA assay (Pierce 23225), then 20ug of each sample was prepared with 6X denaturing loading dye (Boston BioProducts BP-111R) and boiled at 95°C for 5min. Samples were loaded onto 4-12% Bis-Tris gels (Invitrogen NW04120BOX), immersed in MOPS buffer and run at 60V for 10min, 110V for 10min, 160V for 35min, and then transferred onto PVDF membranes (Invitrogen IB24002). Membranes were stained for total protein using REVERT 700 (LiCor 926-11021), imaged and washed, then blocked with 5% dry milk in TBS on a rocker for 1h at room temperature. Primary antibodies (APOE, Invitrogen 701241) were diluted 1:1,000 in 5% BSA in TBS and incubated with membranes on a rocker at 4°C overnight. Membranes were washed three times with 0.1% Tween-20 in TBS (TBS-T), then incubated with HRP-conjugated secondary antibodies (anti-rabbit, Promega W4011) diluted 1:10,000 in 5% milk, on a rocker for 1h at room temperature. Membranes were washed again in TBS-T, then exposed using ECL Prime or Select (Cytiva RPN2232, RPN2235).

### Polysome Profiling

Astrocytes on 10cm dishes were treated with 200ug/mL cycloheximide (Sigma 01810) for 3min and collected with a cell scraper into 500μL lysis buffer (20mM Tris pH 7.5, 150mM NaCl, 5mM MgCl_2_, 1% Triton-X-100, 1mM DTT, 200ug/mL cycloheximide, 1X Halt Protease and Phosphatase Inhibitor Cocktail (Thermo Scientific 78440), 24U/mL DNAse (Promega M6101), 400U/mL RNAsin (Promega N2615)). Lysates were then centrifuged at 13,000g for 10min at 4°C. A 10% sucrose solution (10% sucrose w/v, 20mM Tris, 150mM NaCl, 5mM MgCl_2_, 100ug/mL cycloheximide, 8% glycerol) was layered over a 50% sucrose solution (50% sucrose w/v, 20mM Tris, 150mM NaCl, 5mM MgCl_2_, 100ug/mL cycloheximide, 8% glycerol) and made into a gradient using the BioComp Instruments Model 153 Gradient Station. The lysate supernatant was run through this sucrose gradient using a Beckmann Coulter SW41Ti swinging bucket rotor at 36,000 rpm for 2 hours 45 mins at 4°C. Gradients were fractionated using the Gradient Station and absorbance at 260nm was recorded using a BioComp Instruments Triax Flow Cell. Profiles were aligned using the monosome peak and normalized to the area under the curve of the monosome peak to account for differences in loading.

### Statistical Analysis

All data were organized in Excel then analyzed in GraphPad Prism 11.0.2. Most experiments were analyzed by unpaired t-tests or one-way ANOVAs followed by Tukey’s multiple comparisons tests. Paired t-tests were used for APOE IDP quantification and polysome profiling of BDNF-stimulated astrocytes in Figures 3D and 4H, respectively. For PLA puncta, data for all ACM conditions were collected and analyzed within a single experiment (see Fig S15). Statistical comparisons were performed using one-way ANOVA with Dunnett’s post-hoc correction for multiple comparisons across all four groups. Corrected p-values for each pairwise comparison are reported in the relevant figure panels (Figs. 1E, 3E, 4E).

### Data Visualization

All data were graphed in GraphPad Prism 11.0.2. All bar plots with error bars represent mean ± SEM, all polysome profile line graphs with shaded regions represent mean ± SEM. The following diagrams were created with BioRender.com: Fig. 1A, 2A, 2D, 2F, 2I, 2L, 3A, 3F, 3H, 3J, 3L (left), 3N, 4A, 4F, 5A, 5D, 6A; Supp. Fig. S1A (left), S3A (left), S3B, S4A, S4D, S5A (top), S5B (left), S5D (left), S5F (left), S5G (left), S6D, S8A (left), S9A (left), S10A, S10E (left), S11A (top), S11B (top), S12A (top), S13B, S14A, S14D (left). Figures prepared in Adobe Illustrator 2026.

## ACKNOWLEDGEMENTS

We thank Maggie Mamcarz and Olivia Mosto, as well as Drs. Harrison T Evans, Muhaned S Mohamed, Drew Adler, Shane A Liddelow, Cristina Alberini, and Hyung Don Ryoo for their discussions and technical help. This work was supported by the National Institutes of Health (R35NS122316, EK; T32MH019524, WJL; T32NS086750, WJL).

## AUTHOR CONTRIBUTIONS

WJL and EK designed the experiments and wrote the paper. WJL performed experiments and analyzed data. CCS helped generate primary astrocytes and performed and analyzed assays in neurotoxic reactive astrocytes and treated neurons. MMO performed and analyzed PLA experiments. SVK performed and analyzed polysome profiles. CCS, CJS, EAK, XZ, and RS performed additional western blots, immunocytochemistry, and ELISAs. All authors reviewed and edited the manuscript.

## DECLARATION OF GENERATIVE AI AND AI-ASSISTED TECHNOLOGIES

During the preparation of this work, the authors used Claude by Anthropic during the writing process only to improve the readability and language of the manuscript. The authors reviewed and edited the output as needed and take full responsibility for the content of the published article.

---

**Supplementary Figure S1.**
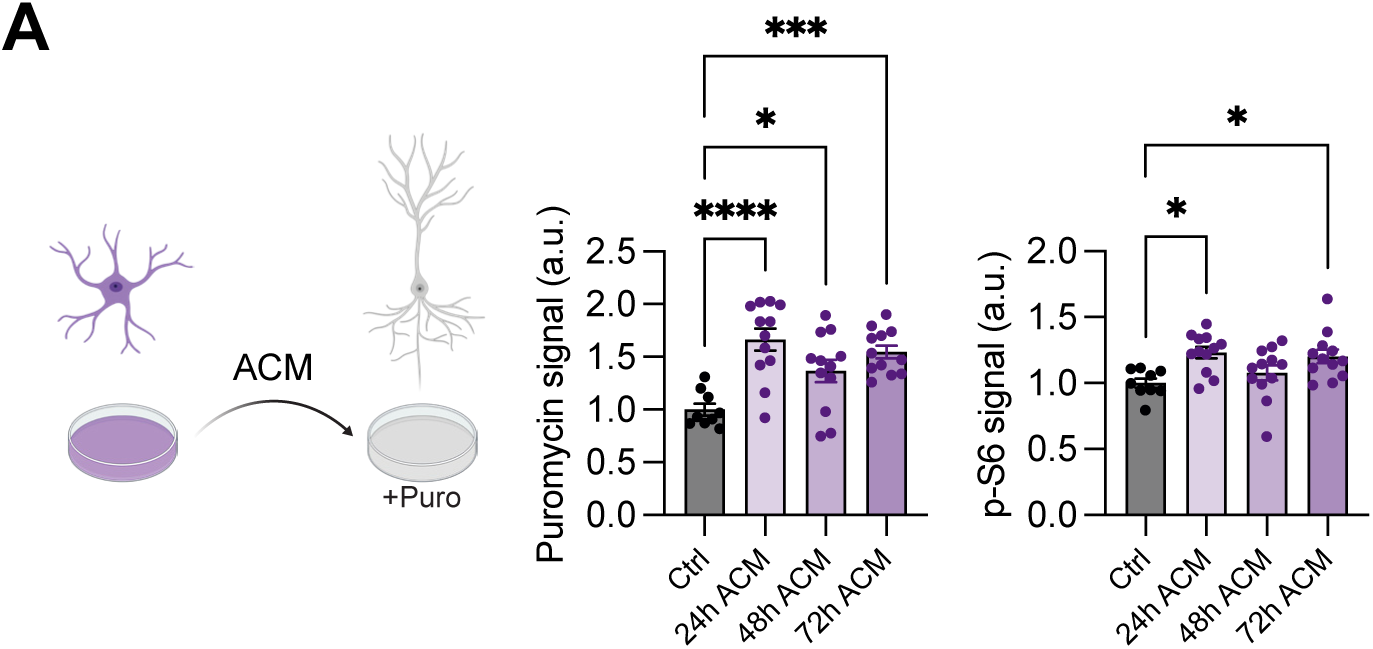
– related to Figure 1B-C. A. Quantification of puromycin and p-S6 signal in MAP2+ neurons treated with ACM collected after 24, 48, or 72h of conditioning. All ACM regardless of conditioning time increase neuronal puromycin and p-S6. Ordinary one-way ANOVA, Tukey’s multiple comparisons test ****p<0.0001, ***p<0.001, *p<0.05, n=3. Bar plots represent mean ± SEM.

**Supplementary Figure S2.**
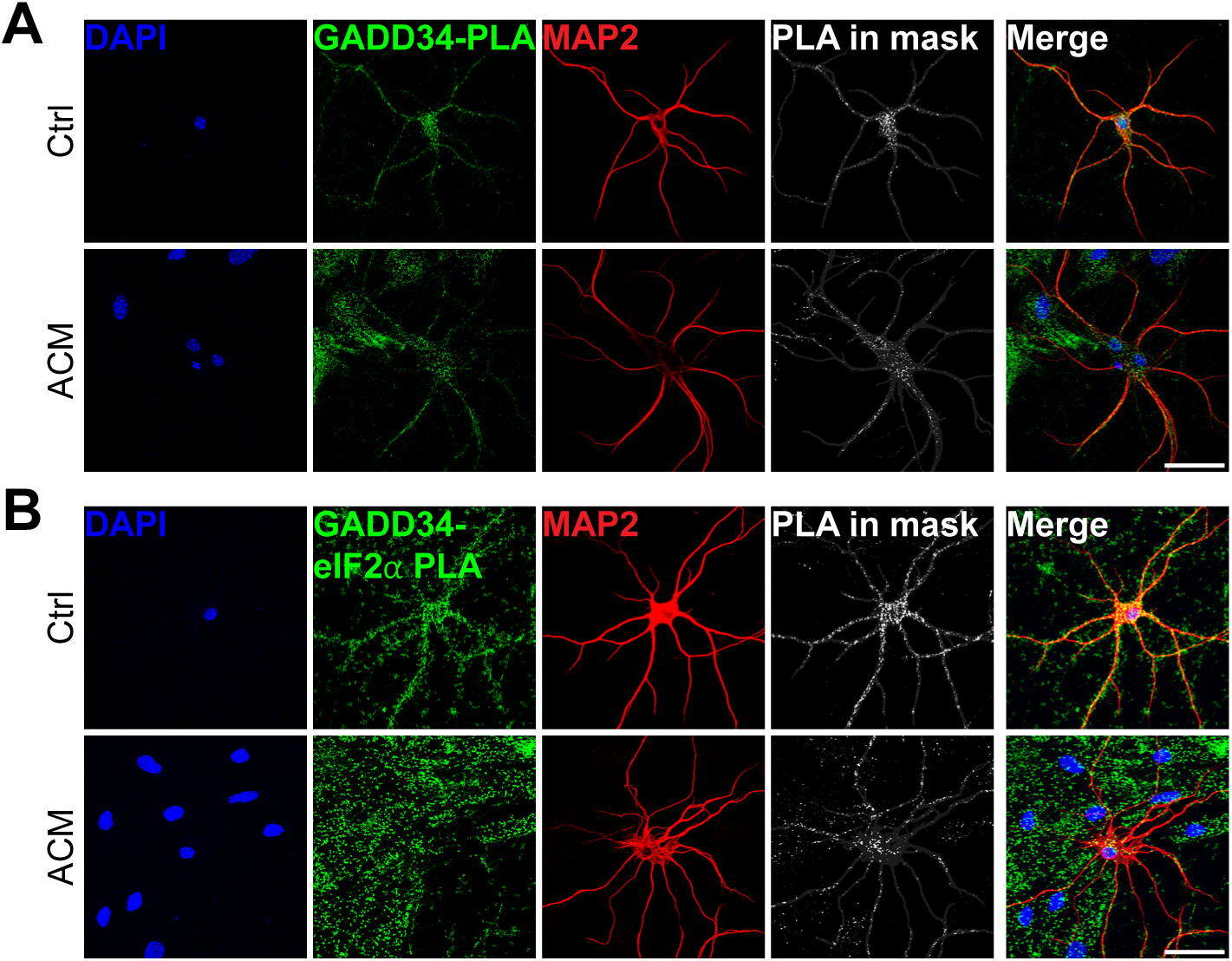
– related to Figure 1D-E. A. Representative images with fully separated color channels, of single protein GADD34 PLA signal in MAP2+ control and ACM-treated neurons. Scale bar=50um. B. Representative images with fully separated color channels, of GADD34-eIF2*a* PLA signal in MAP2+ control and ACM-treated neurons. Scale bar=50um.

**Supplementary Figure S3.**
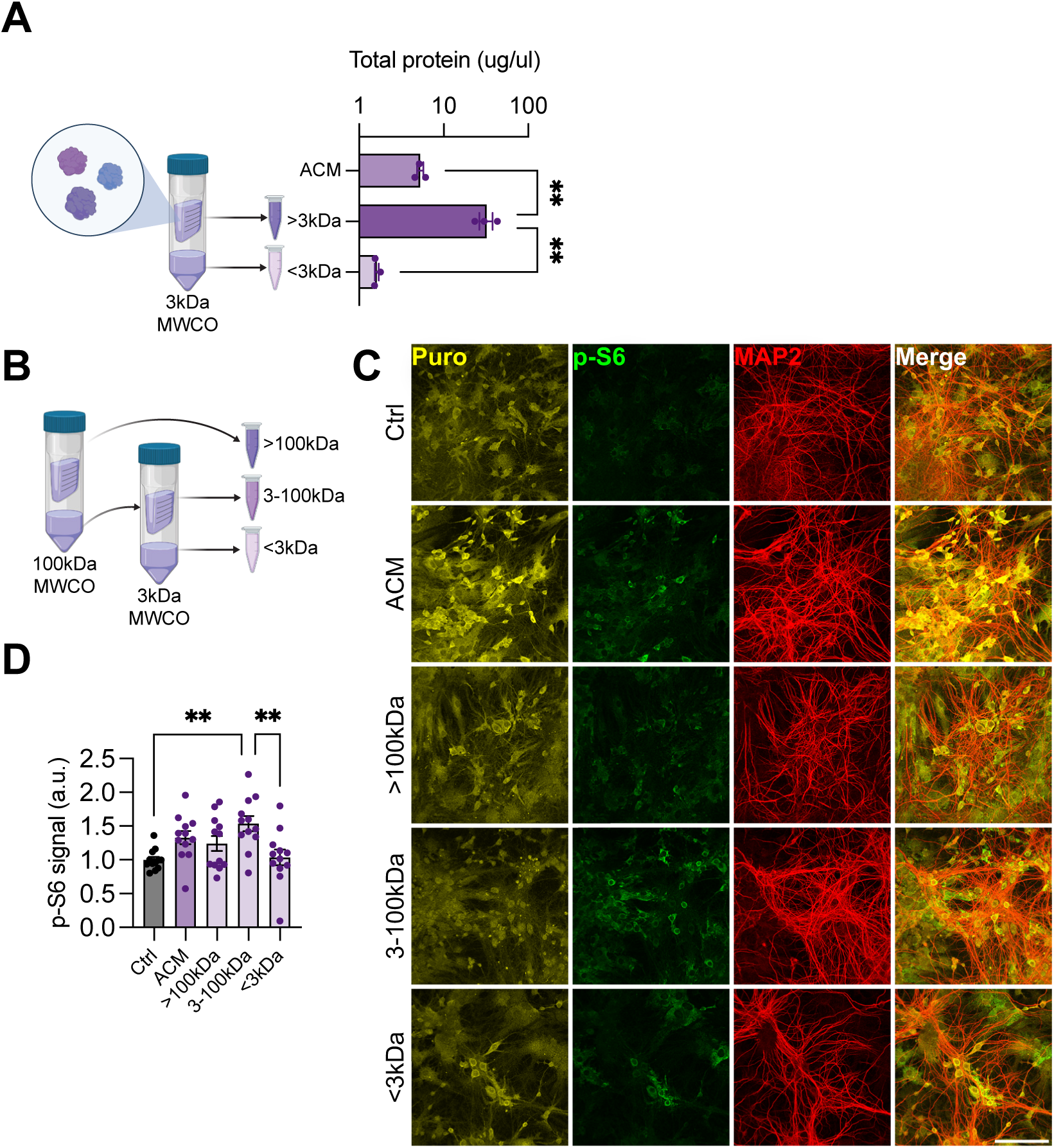
– related to Figure 2A-E. A. Quantification of protein concentration in ACM fractionated through a 3kDa MWCO spin column. The >3kDa concentrate is significantly enriched for proteins. Ordinary one-way ANOVA, Tukey’s multiple comparisons test **p<0.01, n=3. Bar plots represent mean ± SEM. B. ACM was sequentially fractionated through 100kDa then 3kDa spin columns, then the resulting three fractions were used to treat neurons. C. Representative images of puromycin incorporation and p-S6 signal in neurons treated with fractionated ACM. Scale bar=100um. D. Quantification of p-S6 signal in neurons as in panel (D). Ordinary one-way ANOVA, Tukey’s multiple comparisons test, **p<0.01, n=4. Bar plots represent mean ± SEM. Quantification of puromycin signal is presented in Figure 2E.

**Supplementary Figure S4.**
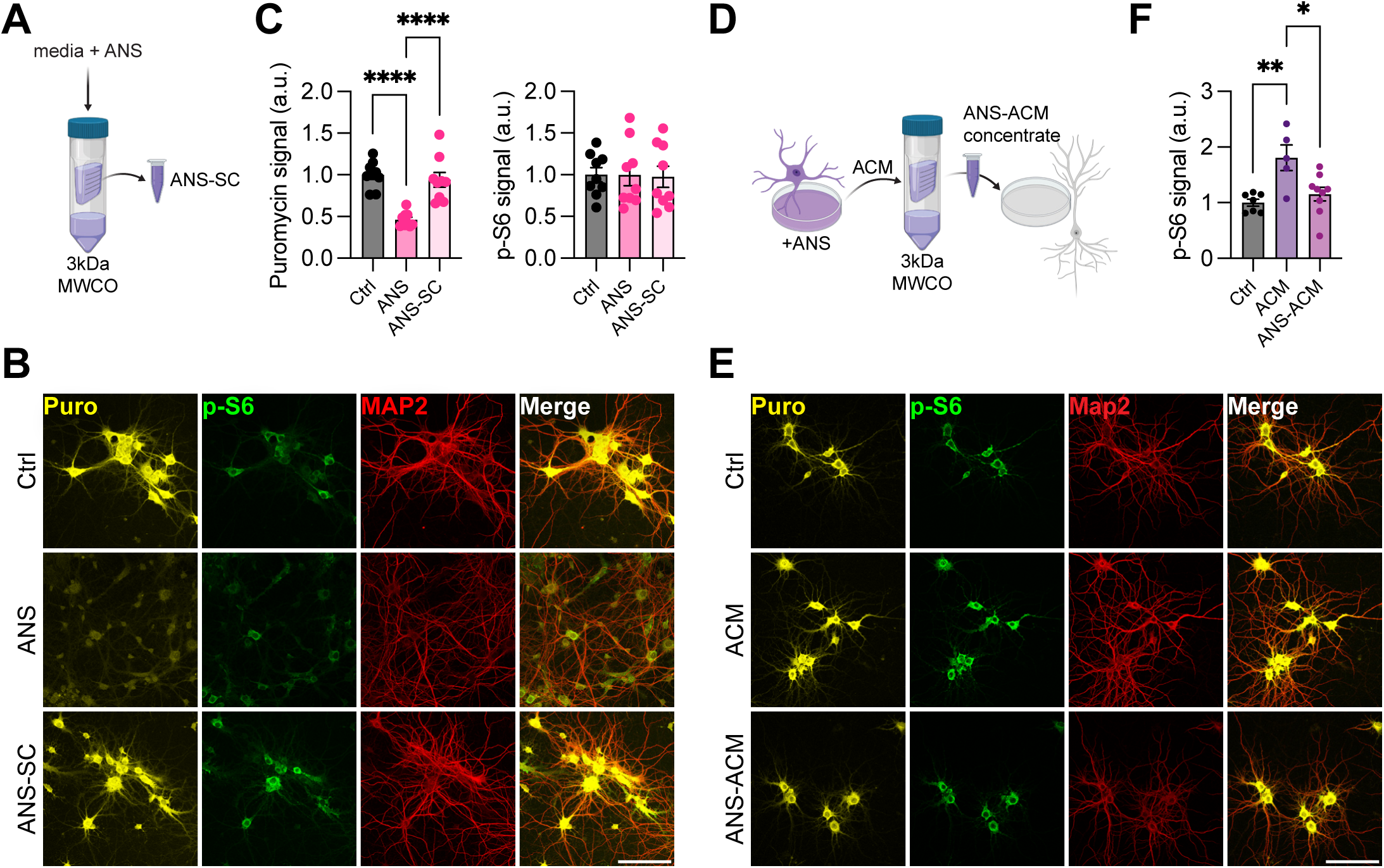
– related to Figure 2F-H. A. Fresh culture medium containing anisomycin was concentrated in a 3kDa MWCO spin column, then used to treat neurons. B. Representative images of puromycin incorporation and p-S6 signal in neurons treated with anisomycin directly (ANS) or with anisomycin filtered through a spin column (ANS-SC). Scale bar=100um. C. Quantification of puromycin and p-S6 signal as in panel (B). ANS-SC does not inhibit neuronal puromycin incorporation. Ordinary one-way ANOVA, Tukey’s multiple comparisons test, ****p<0.0001, n=3. Bar plots represent mean ± SEM. D. Astrocytes were treated with anisomycin (ANS) prior to collecting ACM, whose concentrate was used to treat neurons. E. Representative images of puromycin incorporation and p-S6 signal in control, ACM, or ANS-ACM treated neurons. Scale bar=100um. F. Quantification of p-S6 signal. Ordinary one-way ANOVA, Tukey’s multiple comparisons test, n=3. Bar plots represent mean ± SEM. Quantification of puromycin signal is presented in Figure 2H.

**Supplementary Figure S5.**
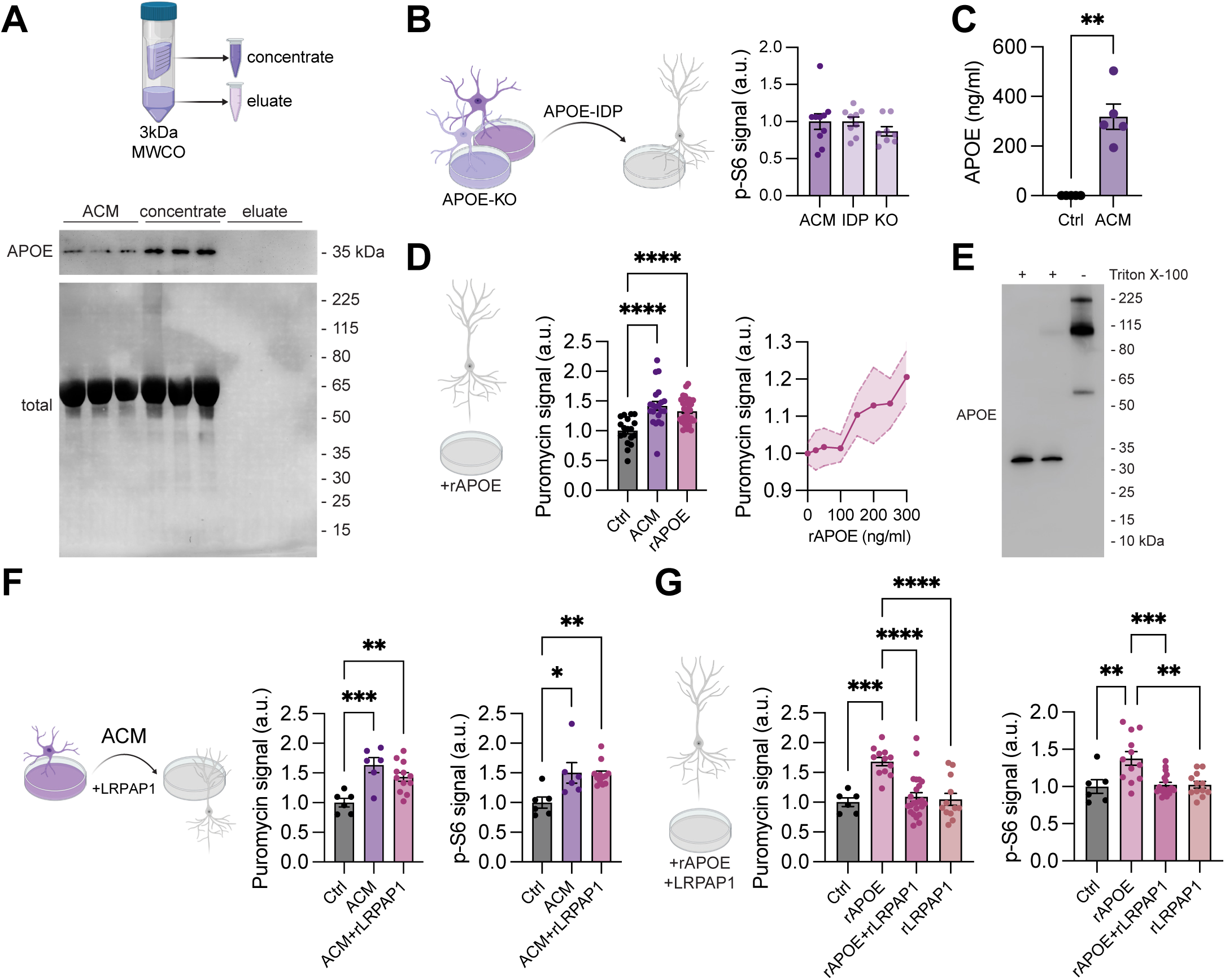
– related to Figure 2I-M. A. Representative western blot of APOE from ACM fractions. APOE is present in the >3kDa concentrate. B. Quantification of p-S6 signal in neurons treated with APOE-IDP or APOE-KO ACM. Ordinary one-way ANOVA, n=3. Bar plots represent mean ± SEM. C. Endogenous APOE concentration in ACM vs. fresh culture medium, detected via ELISA. Student’s unpaired t test, **p<0.01, n=5. Bar plots represent mean ± SEM. D. Recombinant APOE was used to directly treat neurons using 300ng/ml as determined in panel (m). rAPOE increases neuronal puromycin incorporation (left) in a dose-dependent manner (right). Ordinary one-way ANOVA, Tukey’s multiple comparisons test, ****p<0.0001, n=5. Bar plots represent mean ± SEM. E. Western blot of APOE from ACM. APOE only runs at the expected ∼35kDa size with additional detergent, without which heavier bands are observed. F. Quantification of puromycin incorporation and p-S6 signal in neurons treated with ACM alone or with addition of rLRPAP1. rLRPAP1 does not block ACM-driven increase in neuronal puromycin or p-S6 signal. LRPAP1 from two vendors combined. Ordinary one-way ANOVA, Tukey’s multiple comparisons test, ***p<0.001, **p<0.01, *p<0.05, n=2. Bar plots represent mean ± SEM. G. Quantification of puromycin incorporation and p-S6 signal in neurons treated directly with rAPOE alone, rLRPAP1 alone, or both rAPOE and rLRPAP1. rLRPAP1 blocks rAPOE-driven increase in neuronal puromycin and p-S6 signal, with no effect induced by rLRPAP1 alone. LRPAP1 and APOE each from two vendors combined. Ordinary one-way ANOVA, Tukey’s multiple comparisons test, ****p<0.0001, ***p<0.001, **p<0.01, n=2. Bar plots represent mean ± SEM.

**Supplementary Figure S6.**
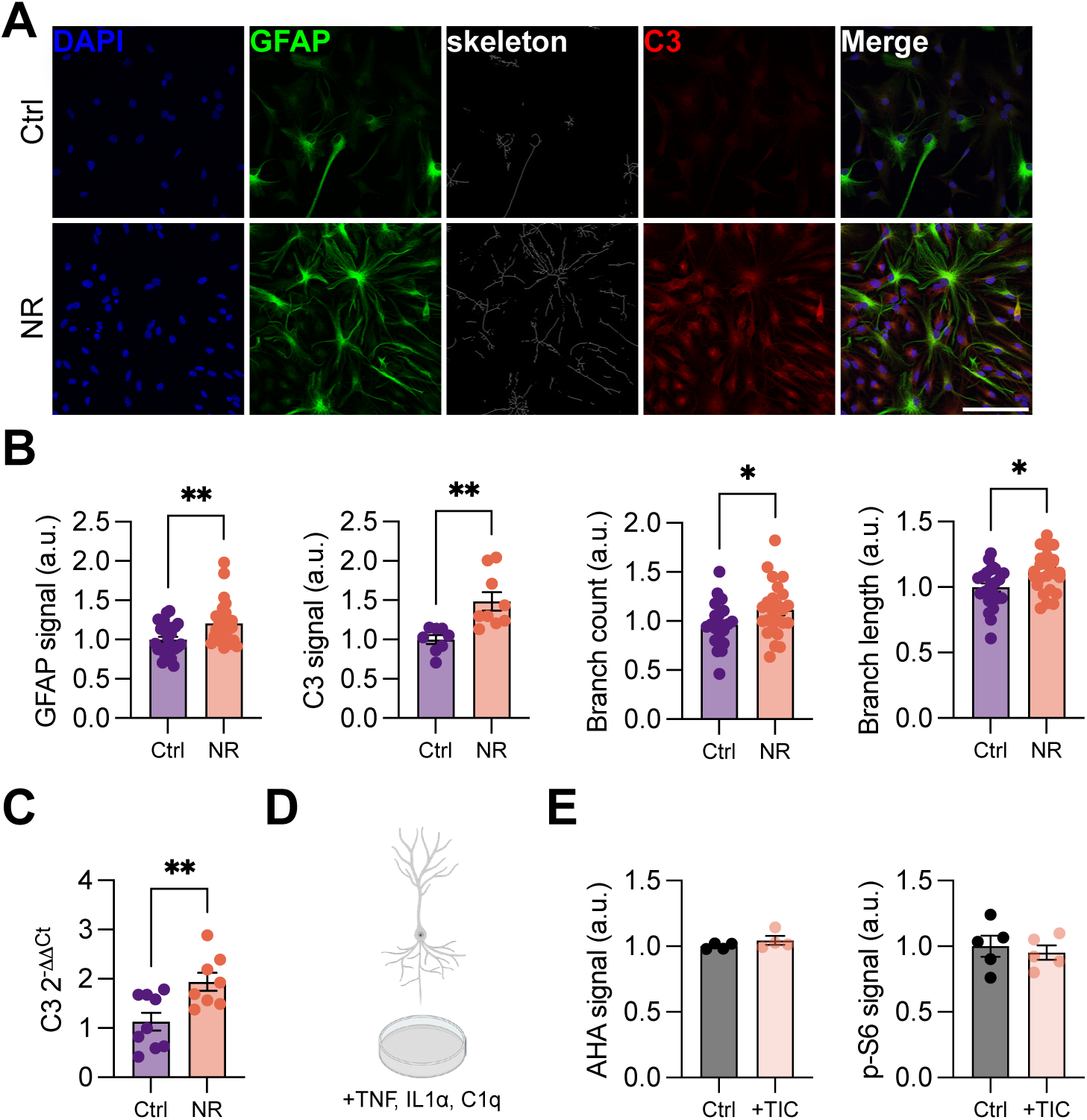
– related to Figure 3. A. Representative images of GFAP and C3 signal in control and neurotoxic reactive astrocytes. Skeleton generated from GFAP signal. Scale bar=100um. B. Quantification of GFAP and C3 signal, and morphological characteristics based on GFAP-skeleton, as in panel (A). NRAs display increased GFAP and C3 signal, as well as increased numbers and length of branches as compared to control astrocytes. Student’s unpaired t tests, **p<0.01, *p<0.05, GFAP n=9, C3 n=3, branch count/length n=22-23. Bar plots represent mean ± SEM. C. NRAs express increased C3 mRNA. Student’s unpaired t test, **p<0.01, Ctrl n=9, NR n=8. Bar plots represent mean ± SEM. D. Neurons were treated directly with TNF, IL1*a*, and C1q. E. Quantification of AHA incorporation and p-S6 signal in neurons as in panel (D). Treated neurons display no change in AHA or p-S6 signal as compared to untreated controls. Student’s unpaired t tests, n=4. Bar plots represent mean ± SEM.

**Supplementary Figure S7.**
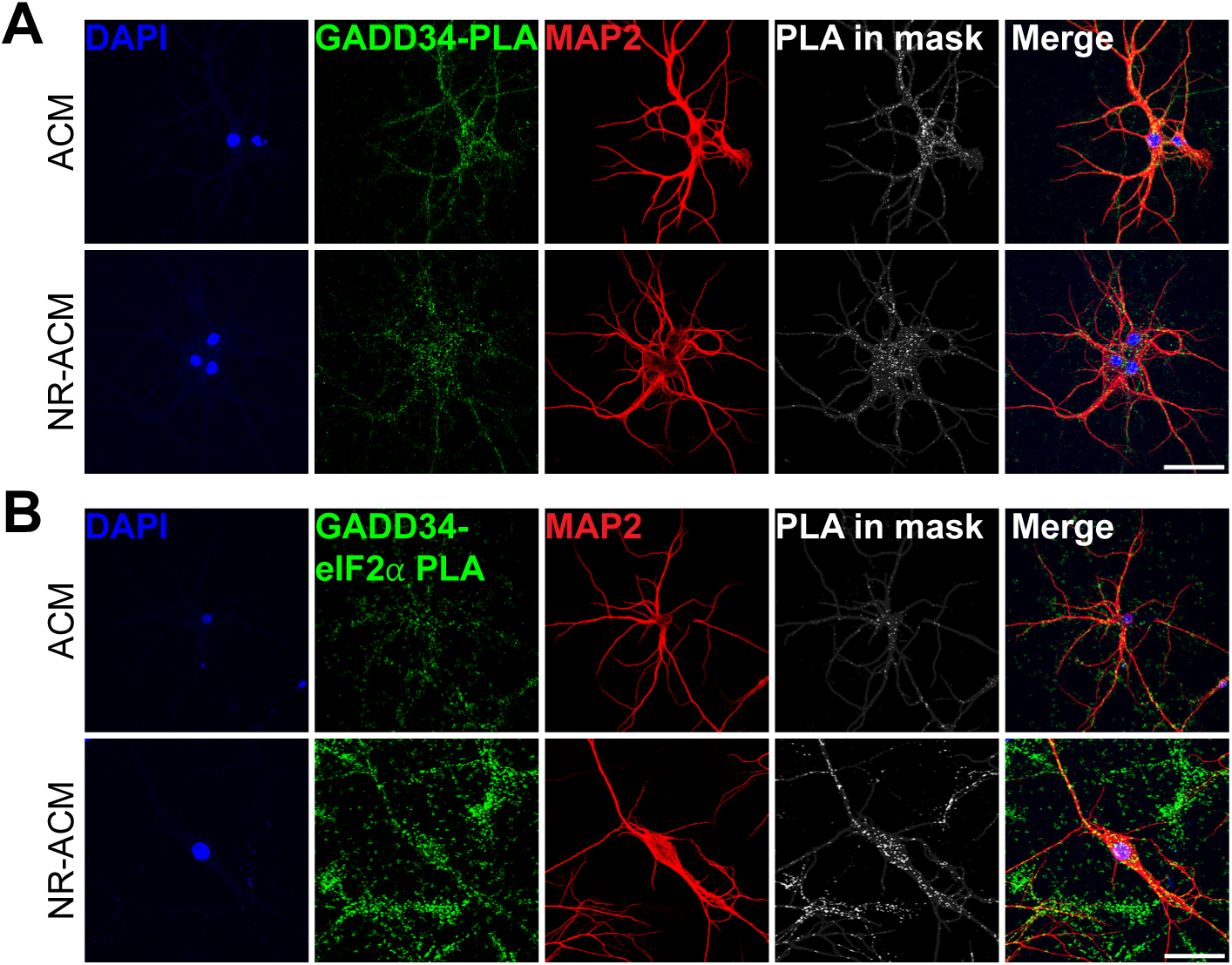
– related to Figure 3D-E. A. Representative images with fully separated color channels, of single protein GADD34 PLA signal in MAP2+ ACM and NR-ACM treated neurons. Scale bar=50um. B. Representative images with fully separated color channels, of GADD34-eIF2*a* PLA signal in MAP2+ ACM and NR-ACM treated neurons. Scale bar=50um.

**Supplementary Figure S8.**
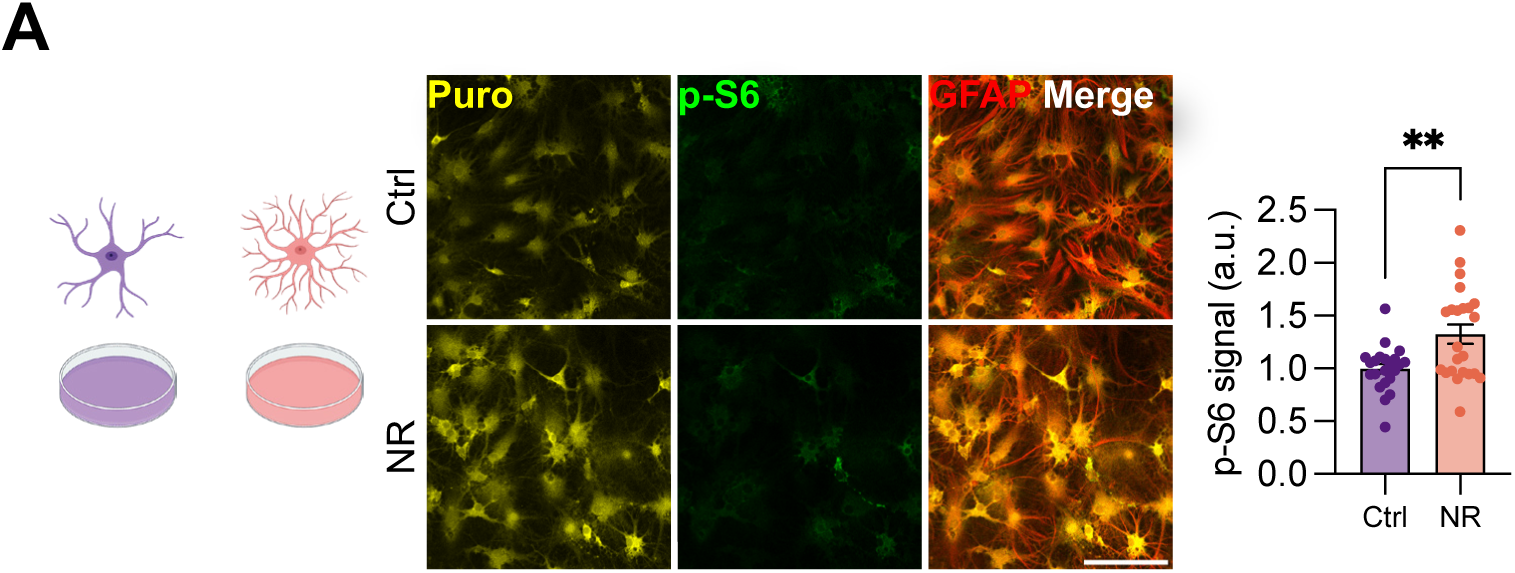
– related to Figure 3O. A. Representative images and quantification of p-S6 signal in NR and control astrocytes. NR astrocytes display increased p-S6 signal as compared to control astrocytes. Student’s unpaired t test **p<0.01, n=6. Bar plots represent mean ± SEM. Quantification of puromycin signal in Fig 3K.

**Supplementary Figure S9.**
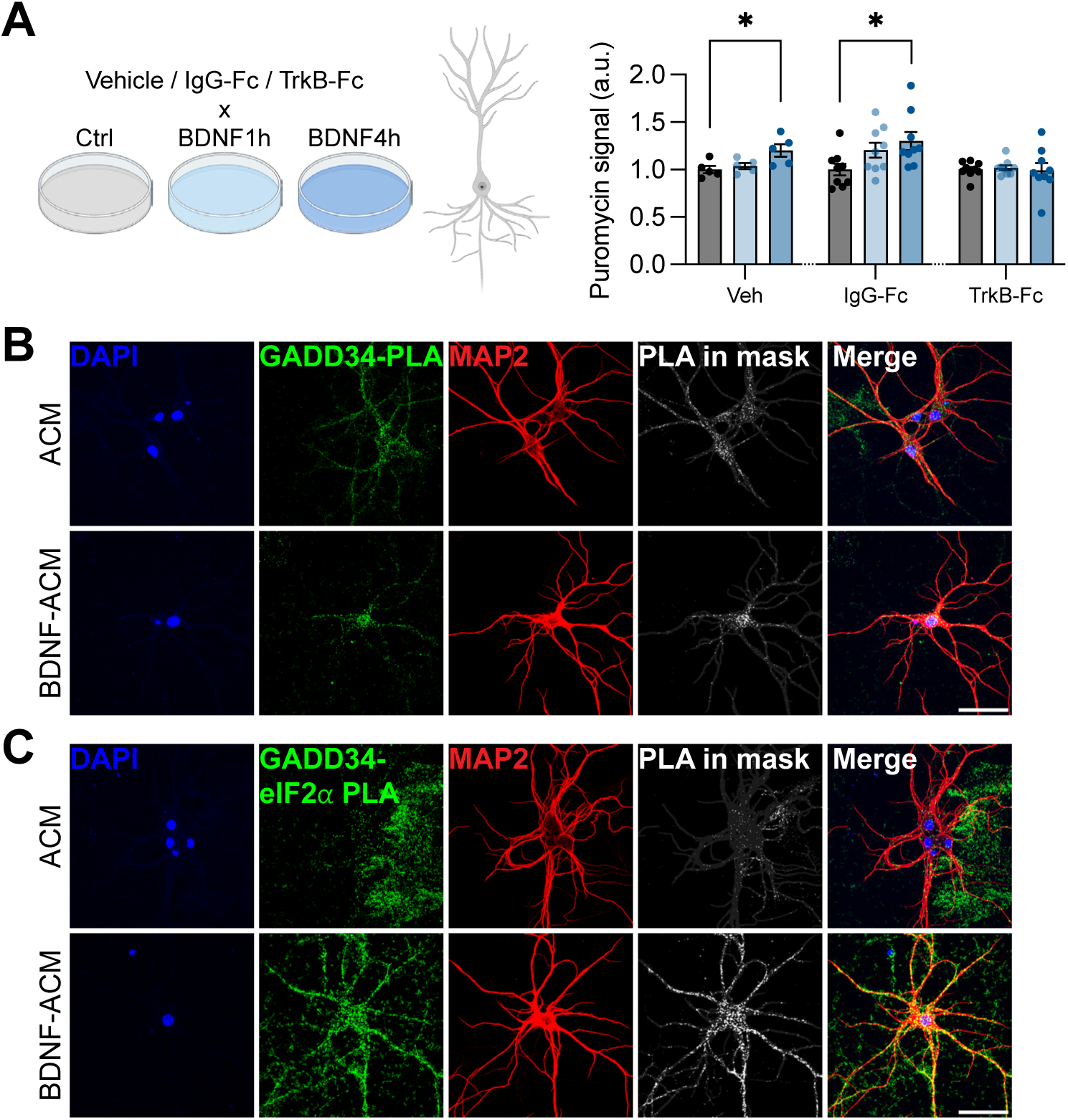
– related to Figure 4A-E. A. Quantification of puromycin incorporation in neurons treated with BDNF for 1h or 4h, and IgG-Fc or TrkB-Fc. TrkB-Fc, but not IgG-Fc, inhibits increased neuronal puromycin signal induced by BDNF activity. Separate ordinary one-way ANOVAs performed for each of Vehicle, IgG-Fc, or TrkB-Fc; Tukey’s multiple comparisons test, *p<0.05, Vehicle n=2, IgG-Fc n=3, TrkB-Fc n=3. Bar plots represent mean ± SEM. B. Representative images with fully separated color channels, of single protein GADD34 PLA signal in MAP2+ ACM and BDNF-ACM treated neurons. Scale bar=50um. C. Representative images with fully separated color channels, of GADD34-eIF2*a* PLA signal in MAP2+ ACM and BDNF-ACM treated neurons. Scale bar=50um.

**Supplementary Figure S10.**
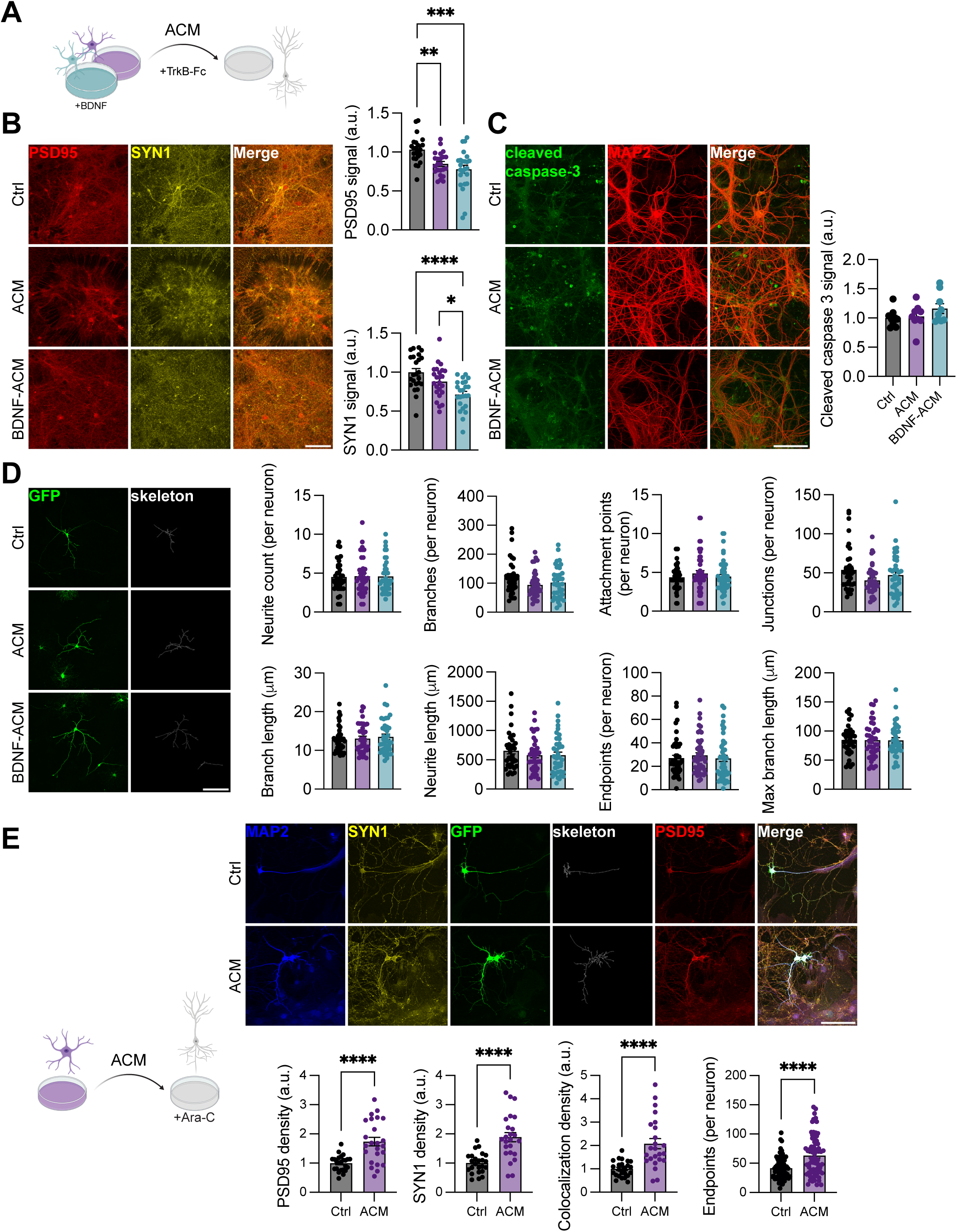
– related to Figure 5B-C. A. ACM, BDNF-ACM, and TrkB-Fc were used to treat neurons. B. Representative images (left) and quantification (right) of PSD95 and SYN1 signal in treated neurons. ACM and BDNF-ACM decreases neuronal PSD95 and SYN1 signal intensity as compared to non-treated controls. Ordinary one-way ANOVA, Tukey’s multiple comparisons test, ****p<0.0001, ***p<0.001, **p<0.01, *p<0.05, n=5. Bar plots represent mean ± SEM. Scale bar=100um. C. Representative images (left) and quantification (right) of cleaved caspase-3 signal in treated neurons. ACM and BDNF-ACM do not change neuronal cleaved caspase-3. Ordinary one-way ANOVA, n=3. Bar plots represent mean ± SEM. Scale bar=100um. D. Representative images (left) and quantification (right) of skeletonized GFP signal in neurons treated with ACM or BDNF-ACM. No gross morphological characteristics change in neurons treated with either type of ACM compared to non-treated controls. Ordinary one-way ANOVAs; Neurite Count n=38-39; Branches n=36-39; Attachment Points n=45-48; Junctions n=37-39; Branch Length n=38-39; Neurite Length n=38-39; Endpoints n=46-47, Max Branch Length n=37-39. Bar plots represent mean ± SEM. Scale bar=100um. E. Top: Representative images of PSD95 and SYN1 signal in neurons cultured with Ara-C, treated with ACM. Neuronal skeletons generated from GFP signal. Scale bar=100um. Bottom: Quantification of PSD95 and SYN1 signal, and endpoint analysis. Pure neuron cultures treated with ACM display increased individual and colocalized PSD95/SYN1 puncta, as well as numbers of endpoints. Student’s unpaired t test, ****p<0.0001; PSD95/SYN1 n=8; Endpoints Ctrl n=65 ACM=70. Bar plots represent mean ± SEM.

**Supplementary Figure S11.**
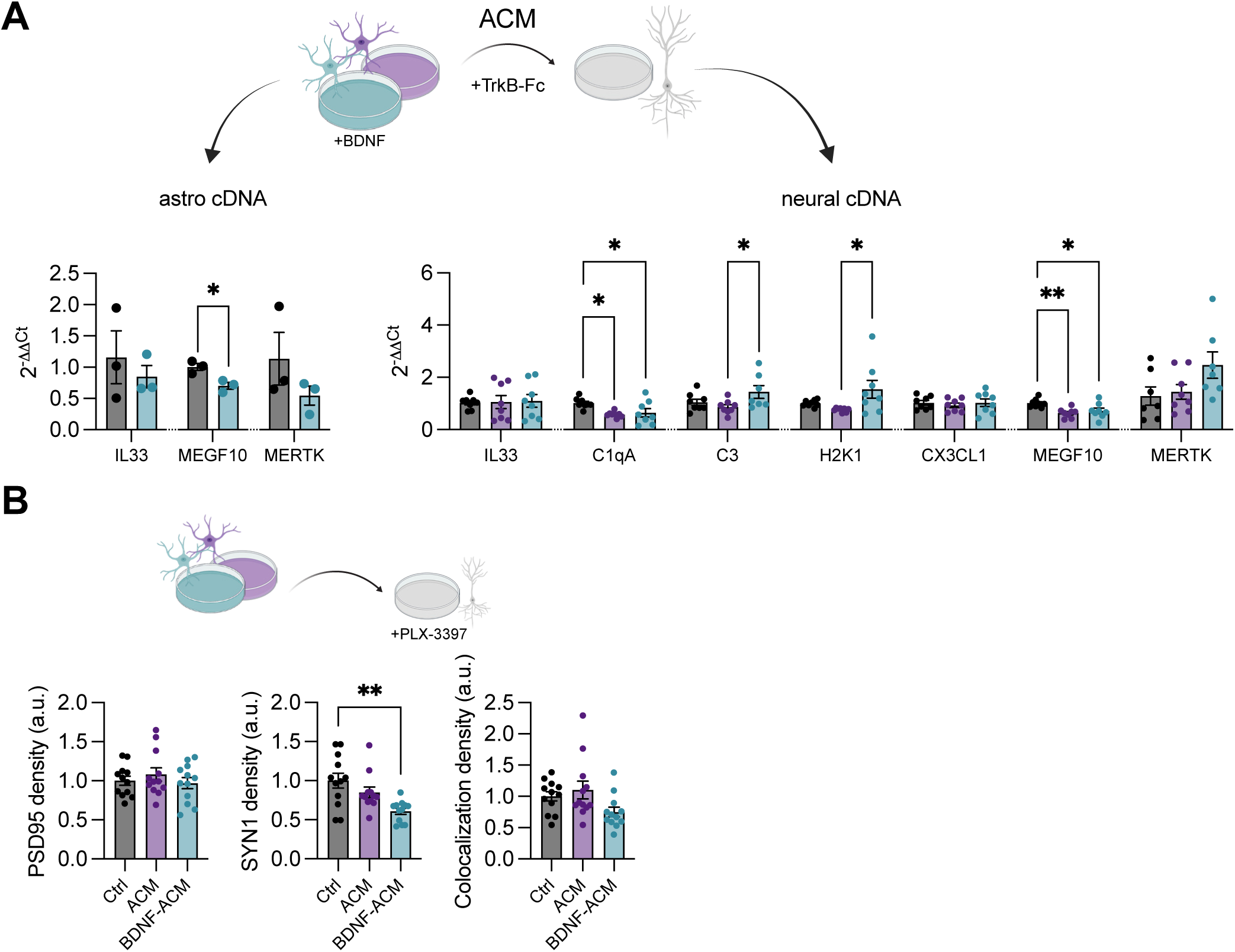
– related to Figure 5B-C. A. qRT-PCR analysis of control and BDNF-stimulated astrocytes, as well as ACM and BDNF-ACM treated neurons. Separate statistical tests performed within each mRNA target. Astro cDNA: Student’s t test, *p<0.05, n=3. Neural cDNA: ordinary one-way ANOVAs, Tukey’s multiple comparisons test, **p<0.01, *p<0.05, n=8. Bar plots represent mean ± SEM. B. Quantification of PSD95 and SYN1 signal in neurons treated with ACM or BDNF-ACM concurrently with PLX-3397. Control neurons with no ACM treatment also received PLX-3397. Ordinary one-way ANOVAs, Tukey’s multiple comparisons test, **p<0.01, n=4. Bar plots represent mean ± SEM.

**Supplementary Figure S12.**
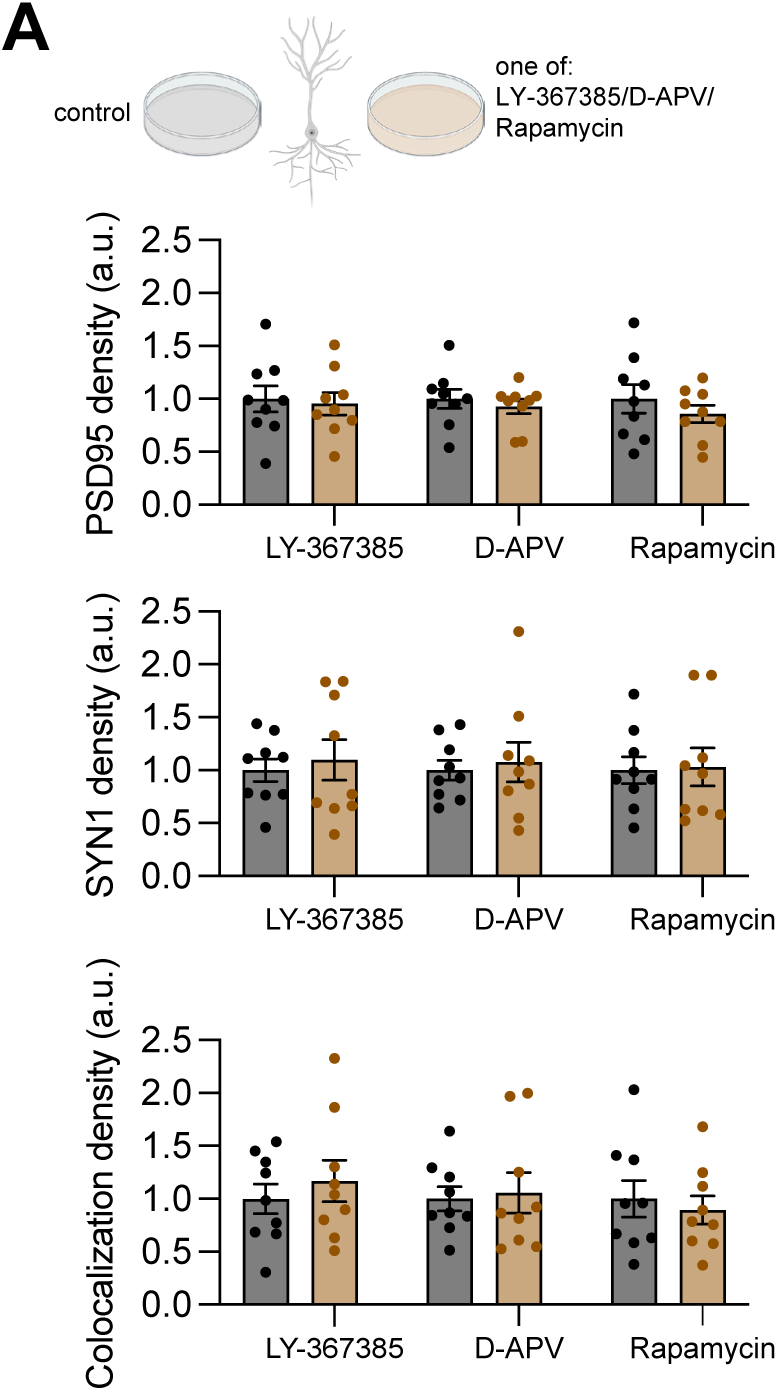
– related to Figure 5D-G. A. Quantification of PSD95 and SYN1 signal in neurons directly treated with one of LY-367385, D-APV, or rapamycin. None of these treatments alter individual or colocalized PSD95/SYN1 puncta at baseline. Separate Student’s unpaired t tests performed within each treatment, n=3. Bar plots represent mean ± SEM.

**Supplementary Figure S13.**
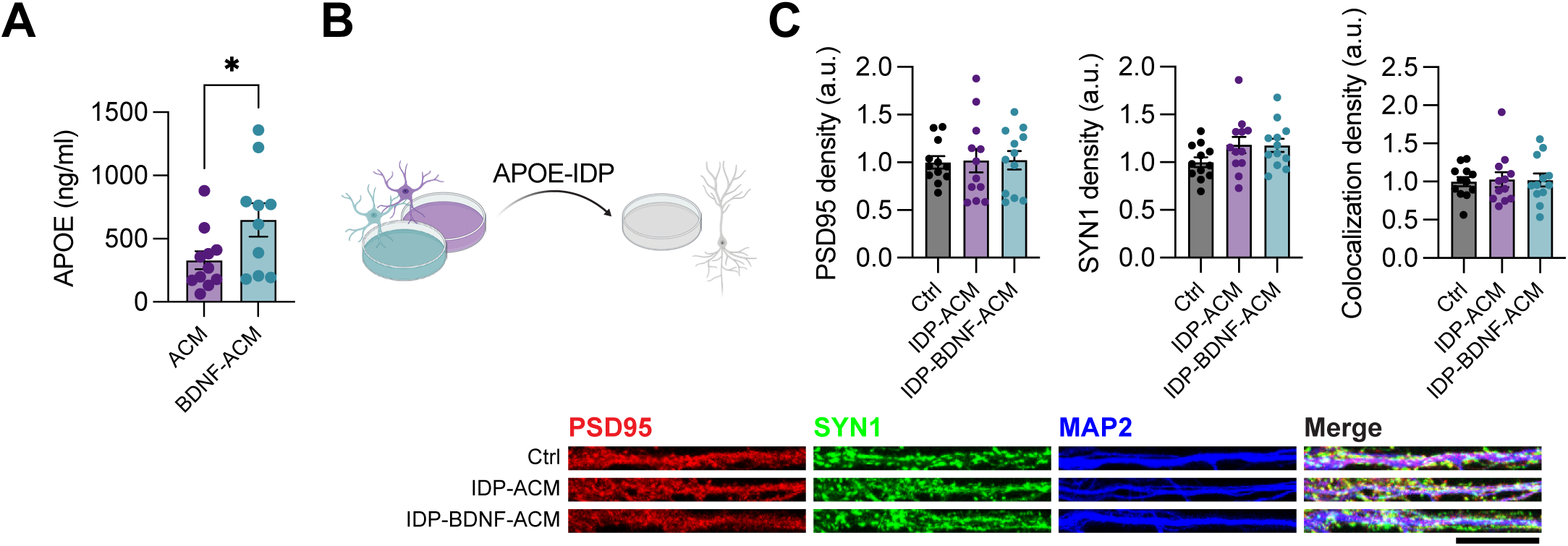
– related to Figure 5. A. APOE concentration is increased in BDNF-ACM vs control ACM, detected via ELISA. Student’s unpaired t test, *p<0.05, ACM n=11, BDNF-ACM n=10. Bar plots represent mean ± SEM. B. APOE immunodepleted from ACM and BDNF-ACM were used to treat neurons. C. Representative images (bottom) and quantification (top) of PSD95 and SYN1 puncta in IDP-ACM treated neurons. APOE IDP blocks the ACM and BDNF-ACM induced decreases in individual and colocalized puncta. Ordinary one-way ANOVA, Tukey’s multiple comparisons test, n=4. Bar plots represent mean ± SEM.

**Supplementary Figure S14.**
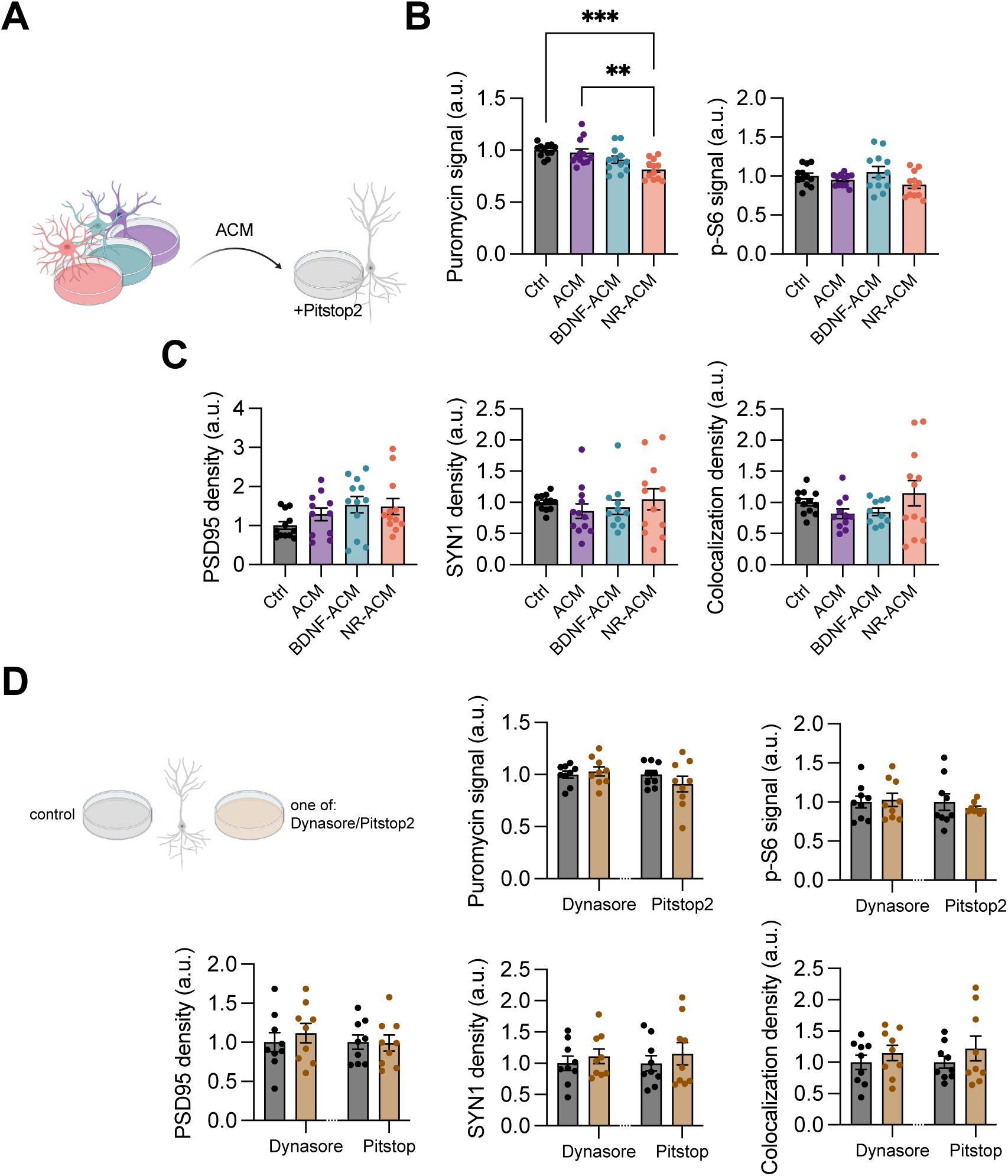
– related to Figure 6. A. Control ACM, BDNF-ACM, and NRACM were used to treat neurons. Neurons were concurrently treated with Pitstop2. B. Quantification of puromycin and p-S6 signal. Pitstop2 blocks increases in neuronal puromycin and p-S6 driven by control ACM and BDNF-ACM. Ordinary one-way ANOVA, Tukey’s multiple comparisons test, ***p<0.001, **p<0.01, n=4. Bar plots represent mean ± SEM. C. Quantification of PSD95 and SYN1 puncta. Pitstop2 blocks decreases in individual and colocalized PSD95/SYN1 puncta driven by control ACM, BDNF-ACM, and NRACM. Ordinary one-way ANOVA, n=4. Bar plots represent mean ± SEM.

**Supplementary Fig S15.**
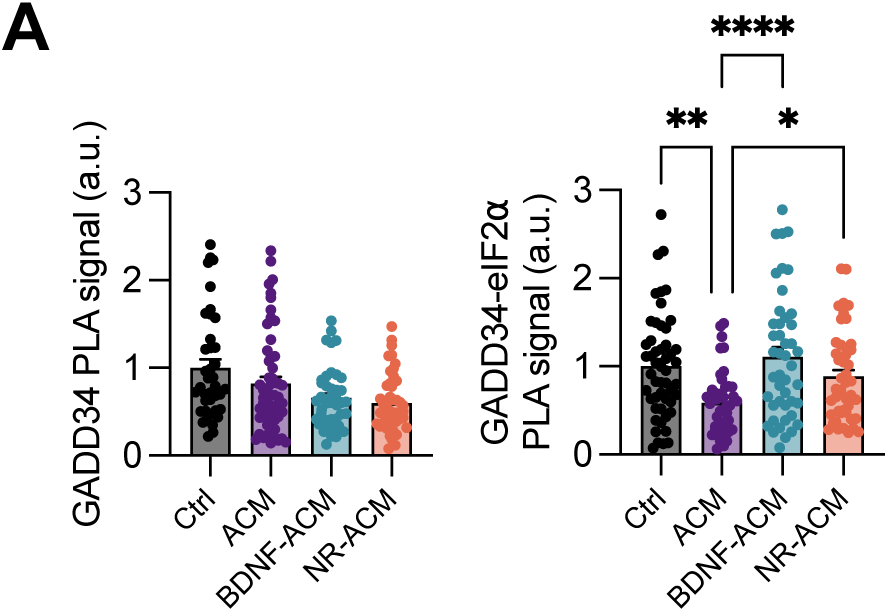
– related to Figure 1E, 3E, 4E. A. Complete dataset of neuronal GADD34 and GADD34-eIF2 PLA interaction across all astrocyte conditioned media conditions. Control, ACM, NR-ACM, and BDNF-ACM treatments were collected and quantified in a single experiment. Data are presented as the complete four-group comparison; statistics were calculated by one-way ANOVA with Dunnett’s post-hoc correction. Corrected pairwise p-values for control vs. ACM, ACM vs. NR-ACM, and ACM vs. BDNF-ACM are reported in the corresponding panels in Figures 1E, 3E, and 4E.

